# Dynamic switching between distinct oscillatory rhythms in prefrontal-amygdala circuits for dimorphic defensive behaviors under natural threats

**DOI:** 10.1101/2022.10.04.510912

**Authors:** Hio-Been Han, Hee-Sup Shin, Yong Jeong, Jisoo Kim, Jee Hyun Choi

**Author notes:** **Contact Info:** Hio-Been Han, Ph.D.,; Hee-Sup Shin, M.D., Ph.D.,; Yong Jeong, M.D., Ph.D.,; Jisoo Kim, M.S.,; Jee Hyun Choi, Ph.D.

## Abstract

The medial prefrontal cortex (mPFC) and basolateral amygdala (BLA) are involved in the regulation of defensive behavior under threat, but their engagement in flexible behavior shifts remains unclear. Here, we report the oscillatory activities of mPFC-BLA circuit in reaction to a naturalistic threat, created by a predatory robot in mice. Specifically, we found dynamic frequency tuning among two different theta rhythms (∼5 or ∼10 Hz) was accompanied by agile changes of two different defensive behaviors (freeze or flight). By analyzing flight trajectories, we also found that high beta (∼30 Hz) is engaged in the top-down process for goal-directed flights and accompanied by a reduction in fast gamma (60–120 Hz, peak near 70 Hz). The elevated beta nested the fast gamma activity by its phase more strongly. Our results suggest that the mPFC-BLA circuit has a potential role in oscillatory gear shifting allowing flexible information routing for behavior switches.

**Highlights:**

- When threatened, mice take quick defensive behaviors such as freeze or flight.
- mPFC-BLA theta tunes its frequency at 5 or 10 Hz for freeze or flight, respectively.
- Low and high theta rhythms in mPFC-BLA emerge in a mutually exclusive way.
- mPFC-driven beta emerges during goal-directed flights, coordinating fast gamma in BLA.

**eTOC Blurb:** Han et al. presents neural dynamics of mPFC-BLA network for freeze-or-flight defensive behaviors under naturalistic threats. Tuning the theta frequency in the mPFC-BLA network is for fast and agile actions under a naturalistic threat, and mPFC-driven beta oscillatory burst is for strategic action.

## Graphical Abstract

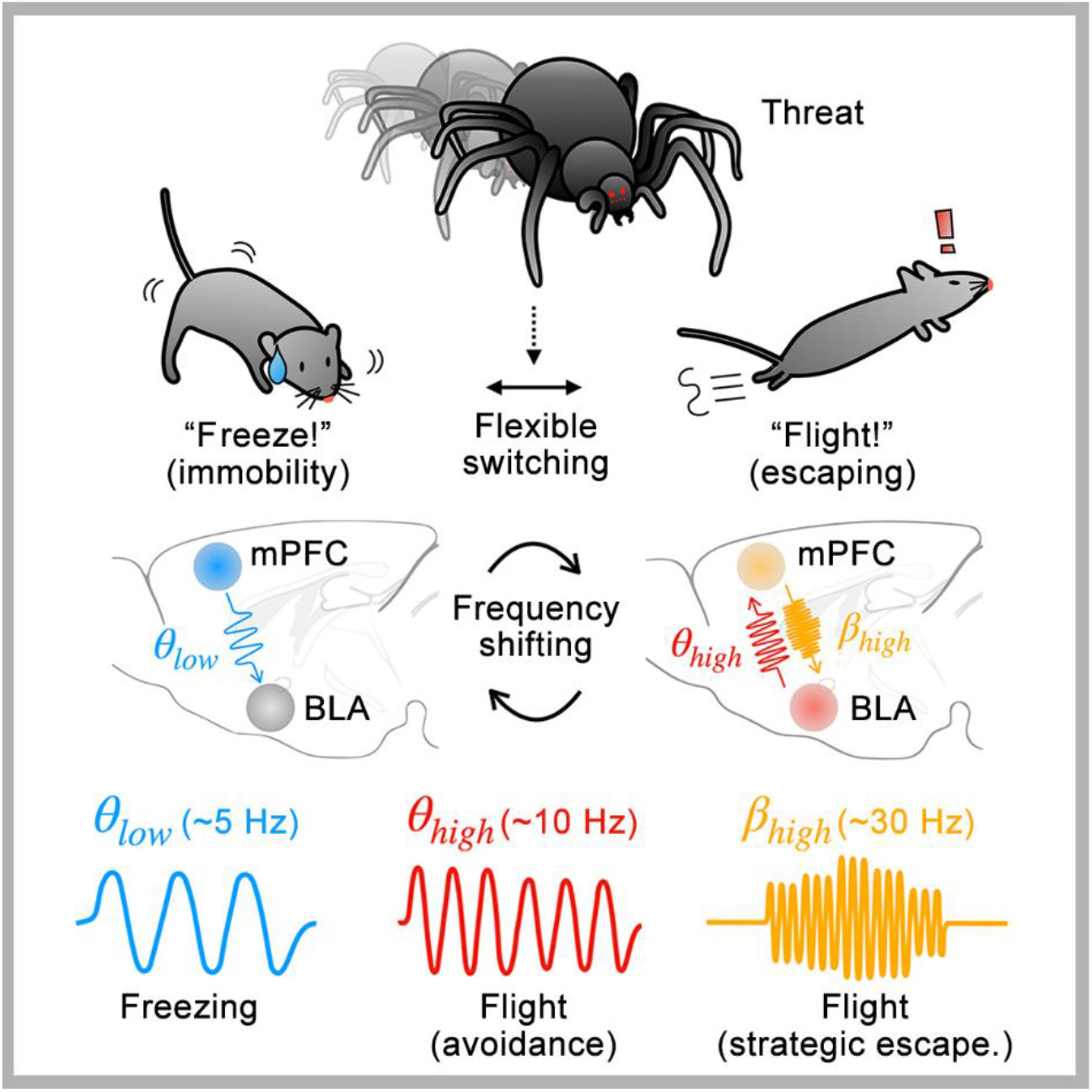

## Introduction

The ability to act appropriately when in danger is crucial for survival. Depending on the context, animals select their defensive behavior in a timely manner—sometimes actively by evading a threat (i.e., flight) or sometimes passively through immobility (i.e., freezing). In the moment of danger, animals are required to instantly assess the level of threat and rapidly exhibit an appropriate behavior followed by continuous interaction with the updated context (Roelofs, 2017). It has been shown that sensory signals guide the selection of behavior (De Franceschi et al., 2016; Hebert et al., 2019), but observational studies of natural behaviors have revealed the complexity underlying contextual determinants (Headley et al., 2019). Moreover, prior experience and/or internal states of the animals can induce certain behaviors (Evans et al., 2019), suggesting that some of the animals’ reactions are the outcome of highly organized information processing in the brain.

In prey animals, an appropriate selection of freezing or flight behaviors under predation risks determines life or death. The freeze-or-flight network in the brain includes the medial prefrontal cortex (mPFC) and the basolateral amygdala (BLA), two areas that are densely and reciprocally interconnected (McGarry and Carter, 2017). Regarding threat-guided defensive action, the mPFC–BLA network integrates threat information with previously encoded associative fear memories to determine viable defense goals (Bukalo et al., 2015; Courtin et al., 2014) and to regulate relevant behaviors through direct projections to the periaqueductal gray (Mobbs et al., 2007; Rozeske et al., 2018) or projections through the central amygdala (CeA) (Tovote et al., 2016). Although the neuronal populations within the mPFC-BLA network engaging in freezing or flight responses have not been fully profiled, this network is known to be involved in shifting between freezing and flight modes (Roelofs, 2017).

One mechanism that long-range interacting neuronal regions use for the flexible routing of information flow is the transient coherence between the activities of the local circuits. A theoretical study showed the emergence of transient synchrony in large-scale circuits mediating flexible routing states and shaping the direction of information flow, reflected by switching patterns of inter-areal phase difference (Palmigiano et al., 2017). Experimentally, evidence has shown that transient coherence between the activities of long-range connected local circuits serves to reorganize the firing phase of local neurons (Benchenane et al., 2010) and to implement attention-dependent signal routing (Grothe et al., 2012; Grothe et al., 2018). Although the inter-areal coherence *in vivo* is stochastic and weak, transient coherence is sufficient to impact information routing and to transition the neuronal state from being irregular to having phase-concentrated activity, which supports the routing-by-synchrony theory (Palmigiano et al., 2017).

While the ethological studies have revealed highly flexible actions of mice within naturalistic environments (De Franceschi et al., 2016; Vale et al., 2017), neural recording of the repertoire of defensive behaviors has been relatively limited and mostly confined to freezing behavior in Pavlovian fear conditioning. Nonetheless, long-range neural synchrony within mPFC–BLA circuits has been revealed to be crucial in freezing behavior (Karalis et al., 2016; Lesting et al., 2011; Stujenske et al., 2014). In particular, the causal role of synchronized activity in the slow theta band (3–7 Hz) in driving freezing behavior has been demonstrated in BLA (Seidenbecher et al., 2003) and mPFC–BLA circuits (Karalis et al., 2016; Lesting et al., 2011; Stujenske et al., 2014). In addition, selective optogenetic manipulation has revealed that mPFC theta oscillation is critical in organizing fear-related functional assemblies at the neuronal level, which leads to successful fear behaviors such as freezing (Dejean et al., 2016). These observations exemplify the communication-through-coherence theory (Akam and Kullmann, 2012; Fries, 2015) and explain the selective neural processing via phase-matching oscillation between the mPFC and BLA during freezing. However, the parallel neural mechanism underlying flight and switching between opposing actions have yet to be identified. In our previous study, we showed that a spider robot as a natural threat could induce activities in the BLA that was recorded by a wireless telemetry (Kim et al., 2020), however neither concurrently measured mPFC nor specific behavioral analysis were not performed.

Here, we have closely evaluated the behavior of individual, freely-moving mice by conducting the frame-by-frame video analysis and investigated the oscillatory dynamics of the mPFC and BLA with respect to the diverse behavioral response to the robot. We hypothesized that the oscillatory activities within mPFC–BLA circuits would mirror the conditions that necessitate the trigger of an appropriate reaction to a threat. Particularly, we tested the prediction that two opposing behaviors, such as freeze-or-flight would elicit distinct oscillatory activity patterns within mPFC–BLA circuits to coordinate behavior state-specific functional ensembles distributed over distant brain regions. We identified distinct frequencies in theta and beta bands within the mPFC–BLA circuits associated with freeze-or-flight defensive behaviors, giving rise to the role of a transient synchrony in flexible behavioral switching. We found that contextual differences shaped the strength and direction of information flow in the mPFC-BLA circuit. Our findings indicate appropriate switching of the neural dynamics in the mPFC–BLA network flexibly routes the flow of information transfer on the fast timescale to support different defensive behaviors in response to naturalistic threats.

## Results

### Mice exhibited two opposite responses to a spider robot - freezing or flight

In our previous study (Kim et al., 2020), mice showed typical defensive responses to a spider robot by naturally responding with freezing or flight. In order to quantify these behaviors, we analyzed neural dynamics in mPFC and BLA as well as concurrent behavioral states from 8 mice (8 sessions per mouse). First, we identified the mouse body using a convolutional neural network (CNN) (see **Methods**), and then the coordinate of the mouse position was determined by calculating the center of mass of the body mask. The recordings were composed of four 1-min stages (consecutive, total of 4 min) as depicted in **Figure 1A**, and we identified the behavioral state of *freeze* and *flight* in the presence of the spider robot (Stages 2 and 3), and the states of *groom* and *explore* in the stages without the robot (Stage 1 and 4). The *flight* and *explore* were automatically identified with kinematic parameters such as acceleration and speed, respectively, and the *freeze* was detected based on the temporal variation in pixel count covering the mouse (i.e., body mask) (see **Method**). The *groom* moments were identified manually in the collected video by the experimenter. The mean percentages of event time are as follows; 10.63% for *freeze*, 9.84% for *flight* under threat presence (Stage 2, 3); 16.56% for *groom*, 8.47% for *explore* under threat absence (Stage 1, 4). All detected events are plotted in **Figure 1B**, and example video clips of behavioral state labeling are in **Movie M1**.

**Figure 1.**
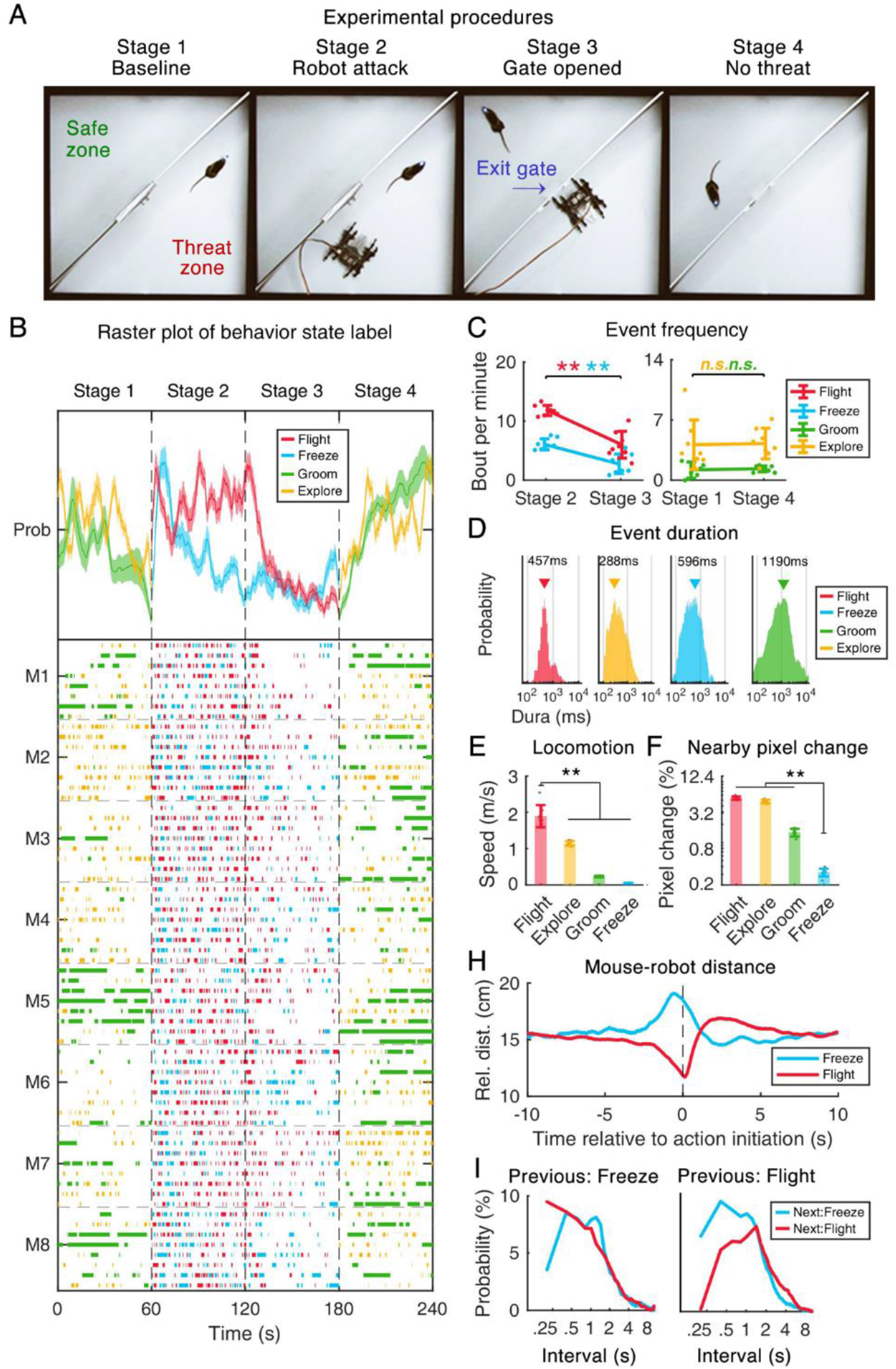
Dimorphic defensive behaviors are induced in the predatory robot paradigm. (A) The experimental paradigm. One session of recording was composed of four one-minute stages (Stage 1: baseline, Stage 2: robot attack, Stage 3: gate opened, and Stage 4: no threat). (B) Raster plot of behavioral states labeled for all mice (M1-M8, n=8) and all trials (trials n=8). Moments of flight and freezing were extracted in the presence of the robot, whereas moments of exploring and grooming were extracted in the absence of the robot. (C) The frequency of behavioral events detected. Gray dots indicate the mean values of individual mice. (D) Histograms of event durations. Upside-down triangle marks indicate the peak value. (E) Comparison of movement speed across behavioral states, showing the fastest speed in flight compared to the speed in other behavioral states. (F) Comparison of the proportion of changed pixels of the mouse body across all behavioral states, showing the smallest value when freezing. Statistical tests were three *post hoc* comparisons (flight vs. others in E, freeze vs. others in F) following one-way ANOVA. (G) Change in robot-mouse distance plotted with respect to the time relative to action initiation at time 0. (H) Time interval between two successive behavioral events. Error bars indicate 1 SEM. ** *p*s < .01.

The occurrence rate shows a sharp increase of *freeze* at the moment of encountering the robot followed by a quick decrease, whereas the occurrence of *flight* did not change over time with the robot (top panel in **Figure 1B**). The occurrence rates of *freeze* and *flight* in Stage 2 were 6.078±0.945/min and 11.813±0.863/min, respectively, and dropped 2.781±1.651/min and 6.031±2.266/min in Stage 3, respectively stage (**Figure 1C**). On the other hand, the total time and occurrence *groom* and *explore* did not change compared to naïve stage (**Figure S1A-B**). While the session repetition did not influence the occurrence in Stage 1 and 2 (**Figure S1C**), *flight* and *explore* in Stage 3 and 4 were reduced with session repetition (**Figure S1D**).

Regarding the behavior features, we analyzed the duration, speed, and pixel changes during each behavior. The behaviors without locomotion, *freeze* and *groom,* and the fast actions, *flight* and *explore* are seemingly similar to each other, but their features were different. For instance, the bout duration of *freeze* was significantly shorter than the duration of *groom* (*freeze* = 0.951±0.092 s, *groom* = 5.567±2.135 s; *post hoc* test, *p* < .001, **Figure 1D**), and the *flight* was significantly faster than *explore* (*flight* = 1.892±0.306 m/s, *explore* = 1.146±0.064 m/s in speed; *post hoc* test, *p* < .001, **Figure 1E**), and the body movement was significantly smaller during *freeze* compared to *groom* (*freeze* = 0.325±0.046 %, *exploration* = 1.472±0.217 % in pixel change; *post hoc* test, *p* < .001, **Figure 1F**). In terms of behavior switch, the mice acted fast enough to change the following behaviors in sub-second levels. For instance, in an adjacent timescale, *flight* is more likely to be followed after *freeze* than after *flight*, and versa (**Figure 1H-I, Figure S2B-C**). This dynamic switch was not path-dependent, that is the choice of action type was not dependent on the type of previous action (**Figure S2D**).

Next, we tested the change of contextual factors that determines the choice of dimorphic *freeze* or *flight* behaviors. As expected, the mouse-robot distance shows that mouse was near the robot at the moment of *flight* and far at the initiation of *freeze* (**Figure 1H**). The analysis on the time interval between two sequential behavioral events revealed switching of action (i.e., *freeze* after *flight*, *flight* after *freeze*) was faster than repeating the same action in sequence (i.e., *freeze* after *freeze*, *flight* after *flight*) (**Figure S2B-C**). However, the choice of action type was not dependent on the type of previous action (**Figure S2D**).

Next, we analyzed the spatial occupancy of mice. Even after the removal of the robot, mice preferred the safe zone to the threat zone, and this preference increased over time (**Figure S2E**), and the latency of exit to the safe zone after gate opening decreased over time (**Figure S2F**). The mice tend to spend more time around the gate area expecting an evacuation during the spider attack period (**Figure S2G**), and spend less time around the corner of the threat zone after the removal of the robot (**Figure S2H**), showing a clear tendency of avoidance of mice regarding the spider robot attack as a harmful event. The level of the threat itself, measured by the proximity of spider-robot distance, was not changed across the sessions (**Figure S2I**).

### Behavior-specific, dimorphic oscillatory states in the mPFC and BLA

To investigate the relationships between the oscillatory activities and behaviors, we analyzed local field potential (LFP) signals in the BLA and the prelimbic area of the mPFC using the CBRAIN wireless headstage (Kim et al., 2020). As shown in **Figure 2A**, the behaviors of mice were rapidly changed during the attack period and the neural dynamics recorded from CBRIAN headstage (**Figure 2B**) showed transient oscillations especially in the theta frequency range. The spectrogram averaged over all 64 sessions showed a prominent increase in oscillatory amplitudes in Stage 2 (**Figure 2C**): during *freeze*, there was one eminent peak in the *θ*_low_ band (3–7 Hz, peak near 5 Hz); during *flight*, there were two prominent peaks, one in the *θ*_high_ band (8–14 Hz, peak near 10 Hz) and one in the *β*_high_ band (22–34 Hz, peak near 28 Hz) (for the bandwidth-of-interest analysis, see **Figure S3A-B** for the baseline noncorrected spectrum). Grand-averaged relative LFP amplitude after the baseline LFP was subtracted showed the same rhythmic behaviors of significantly increased *θ*_low_ activity during *freeze* and significantly increased *θ*_high_ and *β*_high_ rhythms during *flight* (**Figure 2D-E**). The increase in *θ*_low_ during *freezing* was in agreement with previous reports (Karalis et al., 2016; Ozawa et al., 2020), whereas the increase in *θ*_high_ and *β*_high_ of mPFC-BLA circuits during *flight* has not been previously reported. This dimorphic nature of oscillatory activity was not prominent in grooming and exploratory behaviors (**Figure S3C-E**), suggesting that the mere effects of standing still (both for *freeze* and *groom*) or spatial navigation (both for *flight* and *explore*) cannot fully account for these oscillatory patterns during dimorphic defensive behaviors.

**Figure 2.**
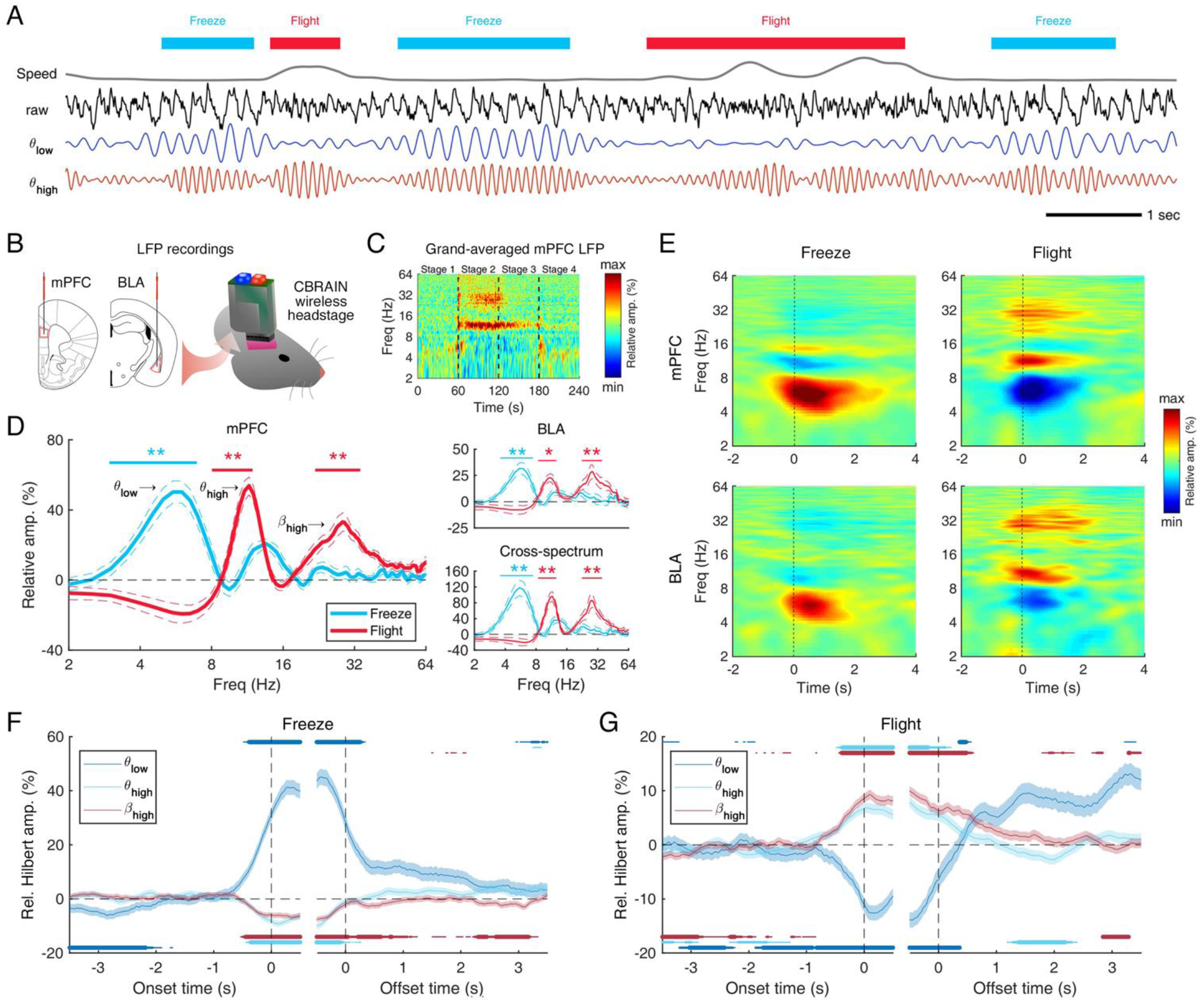
Freezing behavior recruits low theta waves, while flight behavior recruits high theta and high beta waves in the mPFC-BLA network. (A) Example traces of behavioral state, speed, and LFPs recorded from the mPFC of a mouse, showing that *θ*_low_ (3-7 Hz, blue) and *θ*_high_ (8-14 Hz, red) waves emerge alternatively based on the behavioral states of freezing and flight, respectively. (B) Schematic illustration of *in vivo* electrophysiology recording methods. LFP signals from mPFC and BLA were recorded with tungsten wires (left) and transmitted to recording PC with CBRAIN wireless headstage (right). Tungsten wires were implanted in the mPFC and BLA (left) to record LFP signals and CBRAIN wireless headstage (C) Grand-averaged amplitude spectrogram (total 64 sessions from 8 mice) of mPFC LFPs, showing the emergence of three dominant frequency bands: *θ*_low_ at approximately 5 Hz; *θ*_high_ at approximately 10 Hz, and *β*_high_ at approximately 30 Hz) in the threat period (Stage 2). (D) Mean amplitude spectrum for each behavioral state normalized by mean amplitudes in the baseline period. Solid bars indicate the frequency range used in the statistical tests. (E) Time-frequency representation of LFP changes relative to behavioral state onset in the mPFC (top) and BLA (bottom). (F-G) Instantaneous amplitudes via the Hilbert transform relative to the onset and offset of *freezing* and *flight* behavior. Solid bars indicate the statistically significant increase (upper plot) or decrease (lower plot) against the null hypothesis of zero (Wilcoxon signed rank test, uncorrected). The thickness of the bars indicates the significance level (thick: *p* < .001, mid: *p* < .01, thin: *p* < .05). * *p* < .05, ** *p* < .01 tested by Wilcoxon signed rank test against the null hypothesis that the two medians are equal.

To further investigate the temporal relationships between oscillatory activities and behaviors, we calculated the instantaneous amplitude (envelope) of *θ*_low_ and decreased *θ*_high_ and *β*_high_ rhythms via the Hilbert transform and plotted them with respect to the behavioral initiation and cessation. Statistical analysis showed that significant increases occurred before the start of the behavior and persisted after the behavior ended for both *freeze* (**Figure 2F**) and *flight* (**Figure 2G**). To test whether the power of oscillation drives the behavior duration or vice versa, we performed a statistical test regarding the slope in the linear regression to predict power from bout duration. We found positive correlations between *θ*_low_ and *freeze* durations and between *β*_high_ and *flight* durations (**Figure S4**). It is noteworthy that a stronger *θ*_low_ rhythm was found during the longer and more robust (i.e., less movement) *freezing* bouts (**Figure S4C-E**), which replicated the functional property of *freezing*-related *θ*_low_ reported in previous studies (Karalis et al., 2016; Moberly et al., 2018; Ozawa et al., 2020). These results suggest that distinct neural oscillatory mechanisms underlie these two opposing defensive behaviors.

### Mutual exclusivity of oscillatory states in the mPFC and BLA corresponding to freezing and flight behavior

Next, we asked whether these behavior-specific oscillations are exclusive each other and tested the coexistence of oscillations in different frequency bands. We first discretized the spectrogram in binary (see *quantized spectrogram* in our previous work (Han et al., 2017)) by assigning 1 for the significant increase of the power and 0 for the rest. Second, we calculated the conditional probability, p(∃*f_2_* |∃*f_1_*), of having 1 at the target frequency of *f_2_* for a given seed frequency of *f_1_* with a value of 1 across time. Third, the cross-frequency coexistence matrix was constructed by averaging the conditional probability over all sessions (see **Figure S5A** for the detailed calculation procedure). This coexistence matrix captures the co-occurrence between two frequency components within and across regions (**Figure 3A; Figure S5B**). Statistical analysis showed that the occurrence rates of *θ*_high_ and *β*_high_ in the presence of *θ*_low_ were 2.73% and 2.78%, respectively, which were significantly lower than the chance level (5.02%, time-shuffling surrogate, *p*s < .001, Wilcoxon signed rank test; **Figure 3B**). The probability of coexisting *θ*_high_ and *β*_high_ was above the chance level (7.29%), although the difference was not statistically significant (*p* = 0.109). This indicated that although *θ*_high_ and *β*_high_ occasionally emerged during *flight*, the two rhythms did not occur simultaneously and therefore might have distinct behavioral relevance. The calculation of coexistence matrix also captured a strong harmonic component in the theta band, particularly within 5–12 Hz, which may reflect the characteristics of waveform (e.g., sinusoidal shape) in this band.

**Figure 3.**
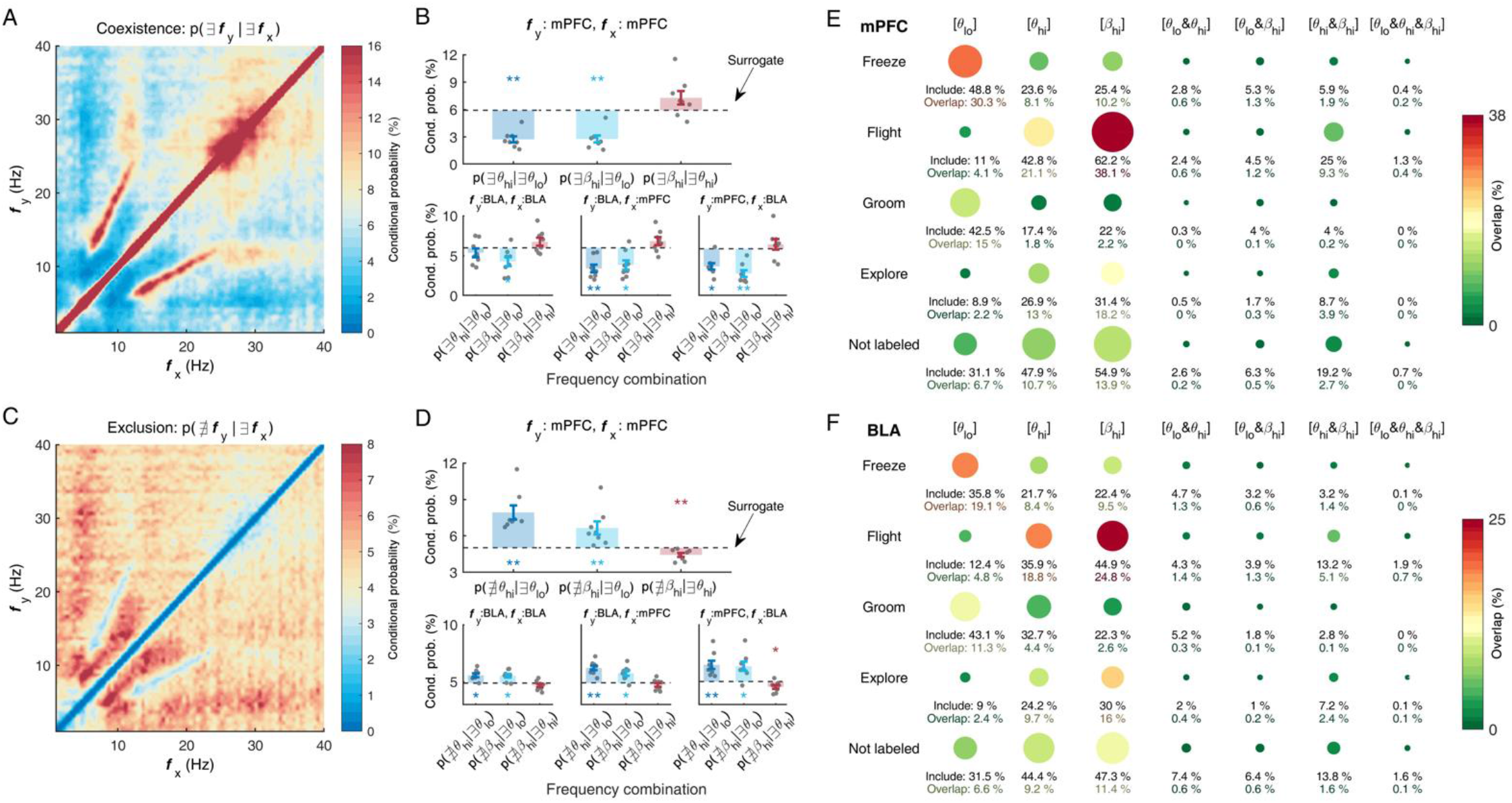
Freeze- and flight-related rhythms are recruited in a mutually exclusive manner. (A) Grand-averaged coexistence matrix obtained from the mPFC of mice, showing the conditional probability of the existence (i.e., top 5% of amplitude) of the frequency component y (i.e., Ǝ*f_y_*) given the existence of another frequency component x (i.e., Ǝ*f_x_*). (B) Mean coexistence probability for three frequency pairs within mPFC (top), within BLA (bottom left), and between mPFC and BLA (bottom middle and right). Black dotted lines indicate the chance levels obtained by surrogate data. (C) Same as (A) but calculated for the exclusion matrix, showing the conditional probability of the nonexistence (i.e., lower 5% of amplitude) of *f_y_* (i.e., ∄*f_y_*) given Ǝ*f_y_*. (D) Same as (B) but for the exclusion matrix. (E) Temporal overlap of behavioral events and oscillatory events. The term ‘inclusion’ indicates the proportion of behavioral bouts that include at least one bout an oscillatory event, and the term ‘overlap’ indicates the proportion of time that behavioral and oscillatory events occur at the same time. Size of the circle indicate the level of inclusion, and color of the circle indicates the level of overlap. (F) Same as (E) but for BLA LFP. * *p* < .05, ** *p* < .01 tested by Wilcoxon signed rank test against the null hypothesis that the median is equal to zero.

Likewise, we calculated the conditional probability, p(∄*f_2_* |∃*f_1_*), of having 0 at the target frequency of *f_2_* for a given seed frequency of *f_1_* with a value of 1 across time and then obtained the exclusion map (**Figure 3C**; **Figure S5C**). The analyses revealed significant exclusivity between *freezing*-related *θ*_low_ and *flight*-related rhythms (all *p*s < .05; **Figure 3D**). The mutual exclusivity of the two theta subbands was particularly interesting, as it provides insight into the underlying circuit dynamics.

### Associated relationship between behavior and rhythms

Next, we asked whether these behavior-specific dynamic patterns are a necessary condition for the correspondent behavior. We calculated the conditional probability for including specific rhythms during behavior and if included, the temporal overlap of two periods of behavior and rhythm. The pie graph in **Figure 3E** shows that approximately half of the *freeze* (48.8%) and *flight* (42.8%) bouts contained at least one *θ*_low_ and *θ*_high_ event in the mPFC. In contrast, *β*_high_ was more often engaged during *flight* (62.2%). Interestingly, neural events of two *flight*-related rhythms that coemerged in the mPFC (i.e., [*θ*_high_ and *β*_high_]) were found in only a quarter of all *flight* episodes (25.0%), signifying their distinct relevance during *flight* behavior. The temporal share of *θ*_low_ during *freezing* was 30.3% in the mPFC, and these values were 21.1% and 38.1% for *θ*_high_ and *β*_high_ during *flight*, respectively. The coemergence of three rhythms (i.e., [*θ*_low_ and *θ*_high_ and *β*_high_]) rarely overlapped with *freeze* (0.2% in the mPFC; 0% in the BLA) and *flight* (0.4% in the mPFC; 0.7% in the BLA) behaviors. These results present the associated but not decisive relationship between the behavior and the behavior-specific rhythms.

### Discontinuous transition of theta frequency

Next, we asked whether the emergence of two theta-rhythmic activities is merely a frequency shift or transition between discrete rhythmic states. Within the range of theta band, it has been reported that frequency shifts in a single brain area can have either *smooth gradients* or *discrete transitions*, possibly through distinct circuit motifs (e.g., chemical changes in a single oscillator, reciprocal inhibition of two oscillators, etc.) (Hernández-Pérez et al., 2016; Vanderwolf, 1969). To characterize the transition pattern, we first collected neighboring *θ*_low_ and *θ*_high_ with a time interval shorter than 3 s (N = 984 for *θ*_low_→*θ*_high_; n = 1014 for *θ*_high_→*θ*_low_). The spectrograms show that both shifts from *θ*_low_ to *θ*_high_ and *θ*_high_ to *θ*_low_ occur in a discrete way rather than in a continuous way (**Figure 4A**). Next, we collected the pairs of two opposite behaviors that occurred within 3 s, and then draw the spectrogram (N = 389 for *freeze*→*flight*; N = 440 for *flight*→*freeze*). Likewise, the frequency transited discretely in both mPFC and BLA (**Figure S6**). These results suggest that the switches between the two theta subbands were discrete rather than continuous.

**Figure 4.**
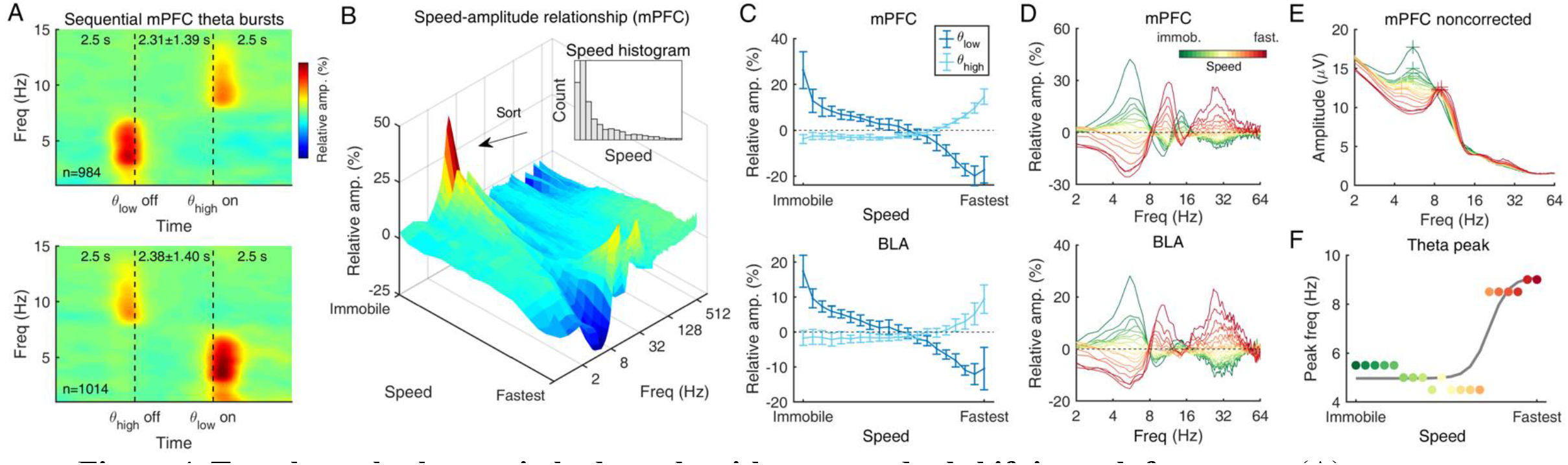
Two theta rhythms switch abruptly without a gradual shift in peak frequency. (A) Time-frequency representation of two sequential theta events, showing a discrete transition from *θ*_low_ to *θ*_high_ (left) and *θ*_high_ to *θ*_low_ (right). Note that data from offset to onset are rescaled to correct for variable time intervals. (B) The amplitude of mPFC LFP as a function of frequency and movement speed. The inset shows an example distribution of movement speed obtained from one session. Note that speed was transformed into rank within individual sessions before binning (bin n=20). (C) Mean amplitude of *θ*_low_ and *θ*_high_ for each speed bin, showing monotonic decrease (*θ*_low_) or increase (*θ*_high_) as a function of movement speed in mPFC (top) and BLA (bottom). (D) Amplitude spectra of the mPFC (top) and BLA (bottom) pseudocolor-coded by movement speed. (E) Baseline non-corrected amplitude spectra in mPFC. Colored crosses indicate the peak in theta frequency range. (F) Change of theta peak frequency detected in (E), showing no sign of middle frequency peak between *θ*_low_ and *θ*_high_. Gray solid line denotes a sigmoidal curve fitting result.

Next, we asked whether the theta dynamics is modulated by engagement of the behavior. We used the speed of locomotion as one-dimensional axis that reflects the behavioral characteristics of both *freeze* and *flight*. The speed-amplitude plot shows two eminent features in the immobile (i.e., a proxy for *freezing*) and the fastest regimes (**Figure 4B**). In particular, the amplitude of *θ*_low_ and *θ*_high_ changed dramatically but continuously along to the speed axis (**Figure 4C**). On the other hand, we did not find any clear evidence of a gradual peak frequency shift (i.e., middle peak around 6–8Hz) in either the mPFC or BLA amplitude spectra across the locomotion speed axis (**Figure 4D**). Regardless of the power, the peaks were located near ∼5 Hz or ∼10 Hz in theta bands (**Figure 4E**), changing in a sigmoidal curve as speed increase (**Figure 4F**). These transition patterns indicate a circuit motif of two distinct oscillators for *θ*_low_ and *θ*_high_.

### Opposite driving directions are present in mPFC-BLA functional networks for freezing and flight

Next, we asked whether the coordination of behavior-specific rhythms exhibit a specific directionality of information flow within the mPFC-BLA circuit. Overall, the LFPs of mPFC and BLA are similar with some similarity (Pearson’s *r* ranged from 0.4 to 0.6), especially under 30 Hz, and this similarity became more obvious when animals were engaged in defensive behavior (**Figure 5A**). Phase locking value (Lachaux et al., 1999) shows that the locking of mPFC and BLA significantly increased at *θ*_low_ during *freezing* and at *θ*_high_ and *β*_high_ during *flight* (**Figure 5B**), implying a potential communication through coherence in these bands. To determine the direction of information flow, we analyzed the lead-lag relationship in the oscillatory networks across different frequency bands by taking the time lag (τ) from the cross-correlation function (exemplified in **Figure 5C**). Replicating previous studies showing that *θ*_low_ during *freezing* in the BLA was driven by the mPFC (Karalis et al., 2016; Ozawa et al., 2020), we found the dominant lead in the mPFC was *θ*_low_. When the locomotion speed was slow (teal line in Figure 6B), the leading pattern in the mPFC became more robust (peak τ = -12.26 ms at 5 Hz; **Figure 5D**). In contrast, directionality was flipped in the *θ*_high_ band, especially when the movement speed was fast (reddish color), with the BLA leading the mPFC at almost zero lag (peak τ = 2.05 ms at 11 Hz). In the *β*_high_ band, the dominant leading direction was from the mPFC, especially when the movement speed was fast (peak τ = -1.51 ms at 32 Hz). The same analyses on the data for *freezing* and *flight* behavioral states revealed a significant mPFC→BLA direction in *θ*_low_ during *freezing* and *β*_high_ during *flight* and BLA→mPFC direction in *θ*_high_ during *flight* (inset in **Figure 5D**). These results emphasize the bi-directional but specific information flow within the mPFC-BLA circuit with respect to *freeze* or *flight*.

**Figure 5.**
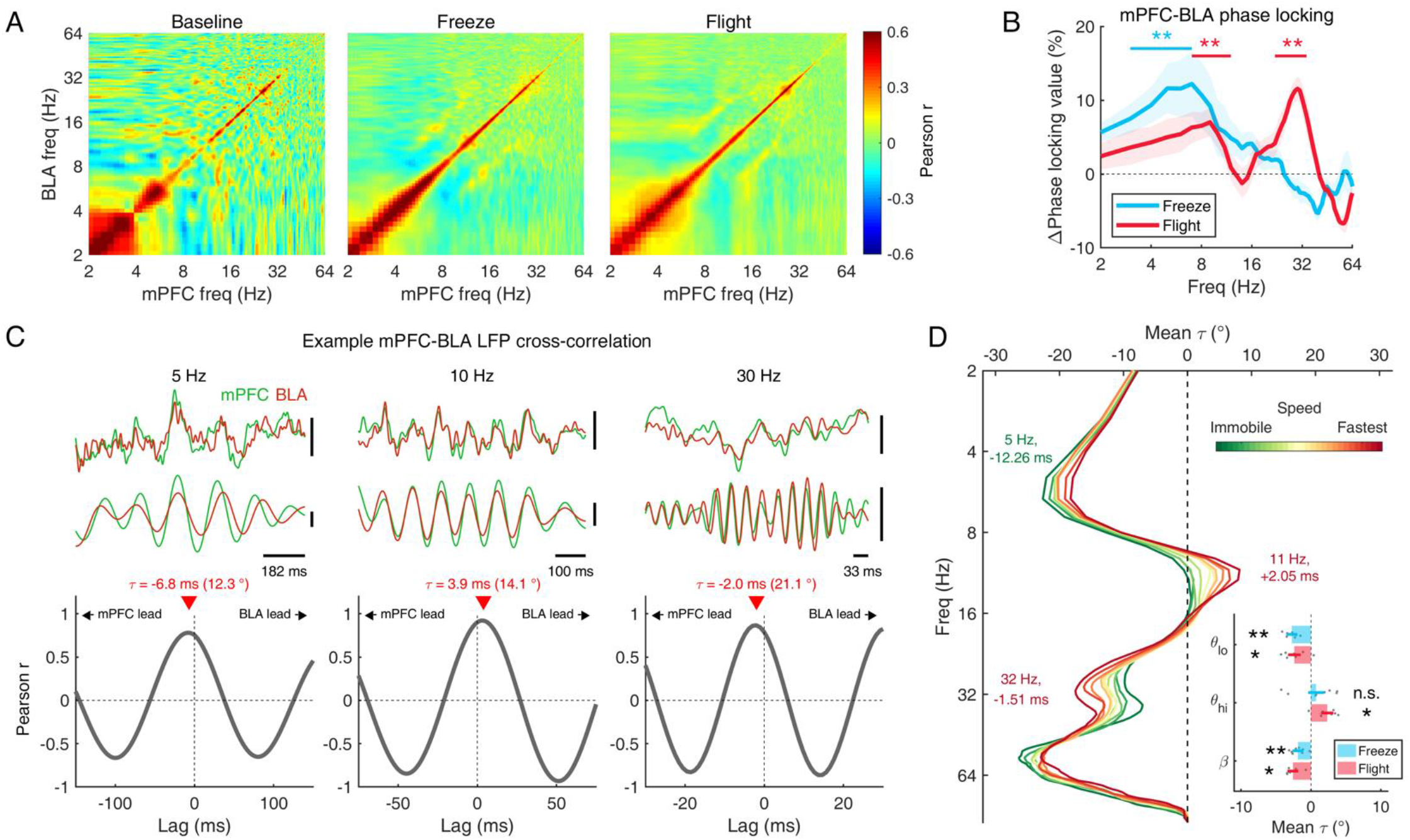
Frequency-specific directionality in mPFC-BLA functional connectivity. (A) Pearson correlations of mPFC and BLA amplitudes showing the clear relationships during the defensive behaviors of freezing (middle) and flight (right) compared to the amplitude in the baseline period (left). (B) Change in phase locking between the mPFC and BLA across the frequency band during defensive behaviors, normalized by the mean of the baseline period. Statistical tests were performed against the null hypothesis that the median is equal to zero (Wilcoxon signed rank test). Solid bars indicate the frequency range used in the statistical test. (C) Examples of mPFC-BLA cross-correlation showing a frequency-specific lead-lag relationship across three frequency components (left: 5 Hz, middle: 10 Hz, and right: 30 Hz). To obtain the cross-correlation function (bottom) in a narrow frequency range, raw LFP data (upper top) were bandpass filtered (upper bottom). The time point of the maximum correlation coefficient, *τ*, was transformed into a phase angle for comparison across a range of frequencies. (D) Mean lead-lag relationship estimated from cross-correlation analysis as a function of frequency (y-axis) and movement speed (color-coded). The text indicates the local maxima of τ in milliseconds. Inset: Comparison of mean τ for each behavioral state in three frequency bands. * *p* < .05, ** *p* < .01 tested by Wilcoxon signed rank test against the null hypothesis that the median is equal to zero.

**Figure 6.**
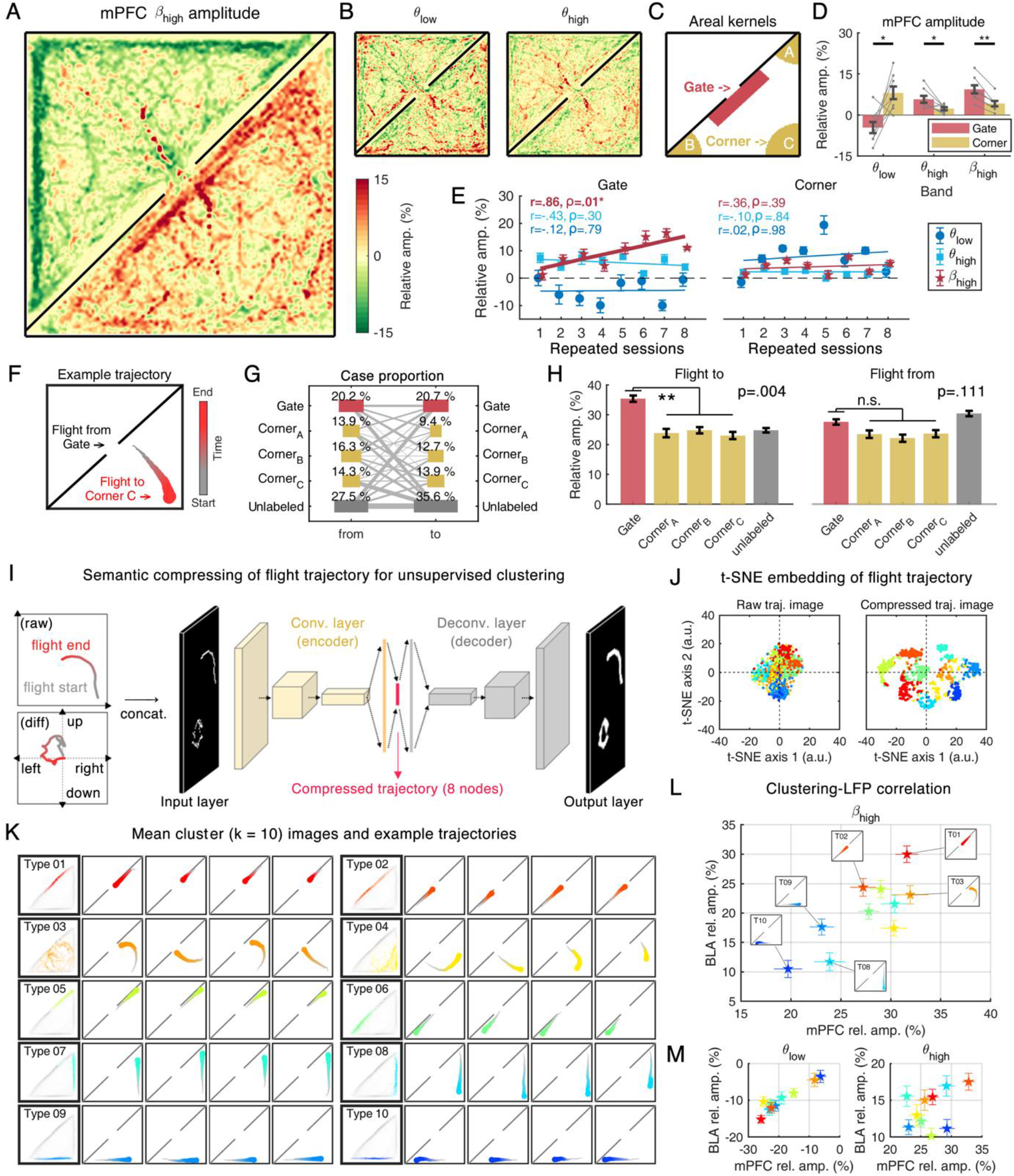
High beta rhythm reflects the top-down cognitive process related to safe-zone-directed flight. (A) Spatial distribution of mPFC *β*_high_ amplitudes from all sessions, showing its spatial selectivity around the gate area in the attack zone. (B) Same as (A) but for other frequency bands. (C) Area kernel masks used for areal mean amplitudes in (D) and (E). (D) Comparison of areal amplitudes. (E) Correlation between repeated sessions and areal amplitudes in each frequency band for the gate (left) and corner (right) areas. (F) Example trajectory of a single flight bout. (G) The proportion of the start (i.e., from) and end (i.e., to) points from all *flight* bouts (bout n=757). (H) Comparison of mPFC high-beta amplitude labeled by the end (left) and start (right) point of *flight* bouts. (I) Schematic illustration of semantic compression for trajectory clustering. Trajectory data (left upper) and its time derivative (left lower) are converted into binary images of spatial occupancy (56*56 each) and compressed in a neural network of a convolutional autoencoder (right). (J) T-distributed stochastic neighbor embedding (t-SNE) of flight trajectory showing a better manifold learning performance after input compression (right) than using a raw trajectory image (left), pseudocolor-coded by the result of k-means clustering (k=10). (K) Mean and example trajectories of each cluster. (L) Cluster-averaged LFP amplitudes during *flight* in the mPFC (x-axis) and BLA (y-axis) in the *β*_high_ band, showing that clusters with gate-directed flights (reddish) tend to include *flight* bouts with a stronger *β*_high,_ while clusters with corner-direct flights (bluish) tend to include bouts with relatively weaker *β*_high_. (M) Same as L, but for *θ*_low_ (left) and *θ*_high_ (right), showing that the relationship of *β*_high_ and clustering results do not appear in other frequency bands. * *p* < .05, ** *p* < .01, *** *p* < .001.

### β*_high_* is associated with goal-directed behavior during flight

We then investigated whether these *freeze-* or *flight-*associated rhythms differed depending on the location or the behavioral patterns. First, we explored the spatial distribution of mPFC rhythms across the arena with a heatmap of the amplitudes relative to the position of the mice that was accumulated across all mice and all sessions. It is noteworthy that *β*_high_ showed a strikingly manifest dependence on the location, while *θ*_low_ and *θ*_high_ did not show such strong location-dependency (**Figure 6A-B**). A closer look showed that *θ*_low_ was stronger when the mouse was in the corner, whereas *θ*_high_ was broadly distributed across the arena but was weaker when the mouse was in the corner. Most importantly, *β*_high_ was dominant around the central gate area. The region of interest (ROI) analysis for the areas near the gate and corner (**Figure 6C**) showed that *flight*-related rhythms, *θ*_high_ and *β*_high_, were significantly stronger around the gate than in the corner, whereas *θ*_low_ showed the opposite pattern (*p* = .234 for *θ*_low_; *p* = .039 for *θ*_high_; *p* = .008 for *β*_high_; Wilcoxon signed rank test; **Figure 6D**). In sum, the *freeze-* or *flight*-related rhythms showed the spatial selectivity to the corner and around the gate, respectively.

Next, we questioned whether this spatial selectivity of rhythms was related to top-down cognitive control, which is enhanced by the past experience. Correlation analyses showed that the amplitude of *β*_high_ around the gate monotonically increased across sessions (all sessions, Spearman’s *ρ* = .86, *p* = .01), whereas no significant correlation was observed in the corner (all sessions, Spearman’s *ρ* = .36, p = .39; **Figure 6E**). In contrast, the two theta rhythms did not show an effect of task repetition (all sessions, Spearman’s |*ρ*|s < .43, *p*s >.30 near the gate; |ρ|s < .10, *p*s >.84 in the corner). However, it was not apparent that the global (i.e., location-invariant) *β*_high_ power during *flight* is increased, as the amplitude of *θ*_low_, *θ*_high_, and *β*_high_ during defensive behaviors in the mPFC-BLA was not changed in the last session compared to that observed in the initial session (**Figure S7**).

Given these observations, we speculated that *β*_high_ would be related to goal-directed behavior (i.e., escape to the safe zone). To test this possibility, we sorted the *flight* bouts based on the positions where the *flight* started and ended with a particular emphasis on the ROIs (**Figure 6F**). Overall, there were no specific relationships between particular start and end positions (**Figure 6G**). However, we found that *β*_high_ in the mPFC was elicited significantly more often during the *flight* bouts heading to the gate compared to bouts heading to the corner (*post hoc p* = .004; Kruskal–Wallis ANOVA, *p* = .006), while the *flight* bouts starting from the gate area did not present such a difference (*post hoc p* = .111; Kruskal–Wallis ANOVA, *p* = .040; **Figure 6H**). As seen in the representative video, the bursts of *β*_high_ are frequently observed in the gate-toward *flight*s, rather than the corner-toward *flight*s (**Movie M2**).

Next, we questioned whether this goal-directed pattern of *β*_high_ can be noticed in the individual *flight*. To deal with the complex spatial patterns during *flight*, we exploited the *flight* trajectories based on their shape (total n = 757). Inspired by unsupervised clustering of handwritten images in machine learning studies (Xie et al., 2016), the 2-dimensional trajectories in position (i.e., absolute position info) and its time derivative (i.e., directional info) were converted to binary images (**Figure 6I**) for unsupervised clustering. To handle the finite number of data samples in our high-dimensional feature space, we compressed the original trajectory image with a dimension of 56 × 56 × 2 into the compressed into a dimension of 8 × 1 using a convolutional autoencoder (see **Methods**), and then a K-means clustering was conducted. To determine the optimal number of clusters, we calculated the goodness of clusters based on maximal separation, and k=10 was determined (**Figure S8A**). Manifold embedding dramatically contrasts the effect of data compression (**Figure 6J**), which enabled successful clustering of the *flight* trajectory (**Figure 6K, Figure S8B-C**). We found that some of the clustered trajectories were instances of gate-directed *flight* (Types 01, 02, and 03 in **Figure 6K**). Likewise, some clusters represented instances of escaping into the corner (Types 08, 09, and 10 in **Figure 6K** and **Figure S8C**).

Next, we calculated the mean amplitude of *θ*_low_, *θ*_high_, and *β*_high_ in each trajectory type. It was noticeable that the gate-directed trajectories were in the high level of *β*_high,_ whereas the corner-directed trajectories were in the low level of *β*_high_ (**Figure 6L**). On the other hand, this discrepancy was not observed in another flight-related rhythm, the *θ*_high_ (**Figure 6M**).

After opening the exit gate to the safe zone in Stage 3, the entrainment of *β*_high_ in mPFC markedly decreased after entering the safe zone (**Figure 7**). The occurrence of *β*_high_ was cut in half after mouse getting into the safe zone (**Figure 7D**). As session repeated for 8 times, the duration of *β*_high_ burst before crossing the gate increased, although the total number of occurence wasn’t changed (**Figure 7E-F**). All these goal-directed patterns suggest a putative role of *β*_high_ in the top-down processes under threat, seeking after the safety guided by previous experience.

**Figure 7.**
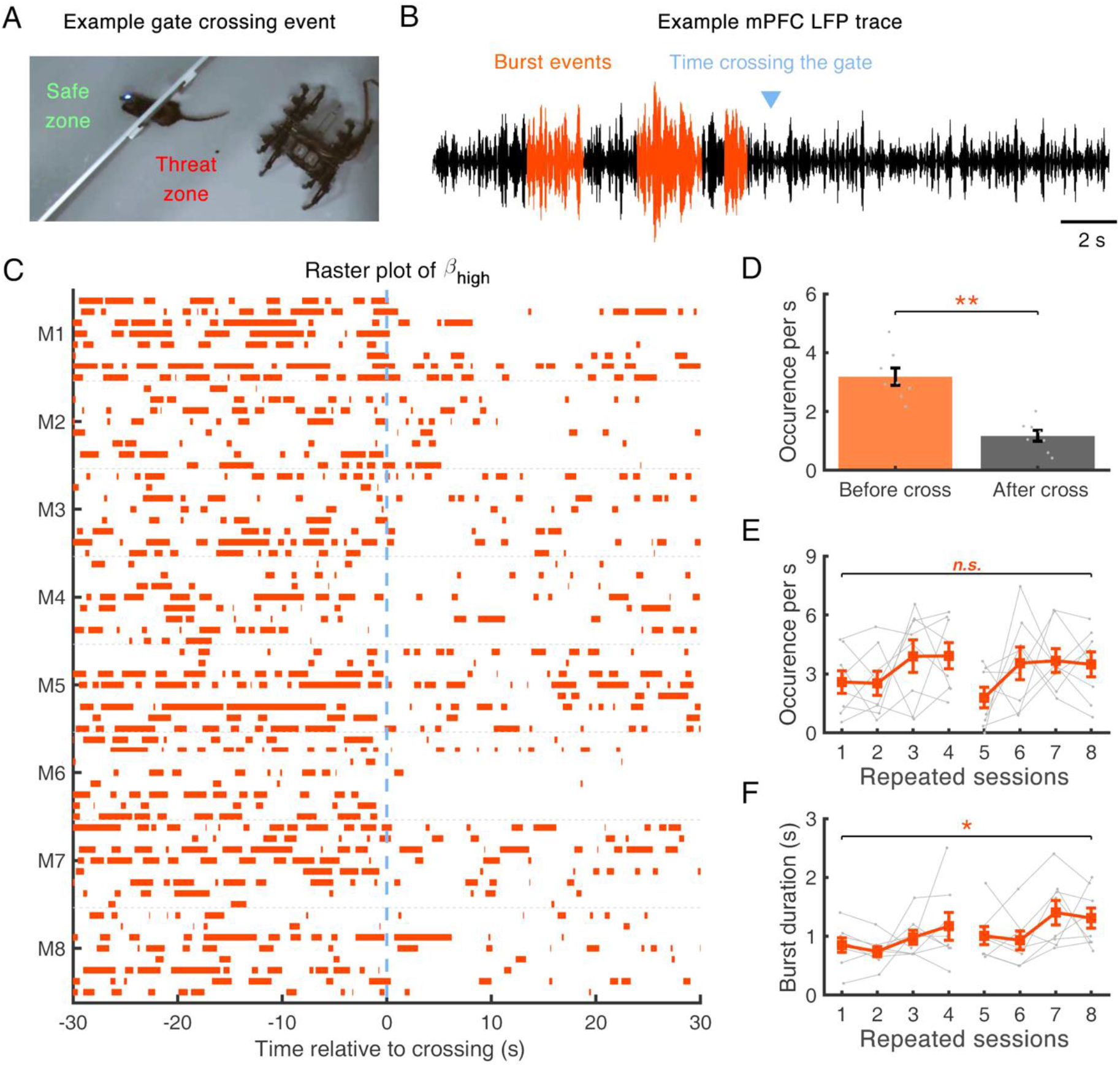
Silencing of high beta activity after a gate-crossing event from the threat zone to the safe zone. (A) Example captured frame of the gate-crossing moment. Note that the gate-crossing moment was defined as the first frame in which half of the body had crossed into the safe zone during Stage 3. (B) Example LFP trace filtered in the *β*_high_ band. Detected burst periods are marked in orange. (C) Raster plot of *β*_high_ bursts, time relative to gate crossing, which occurs at 0 sec. (D) Comparison of the occurrence rate of *β*_high_ bursts before (left) and after (right) animals crossed the gate (Wilcoxon signed rank test, *p* < .01). Burst events were detected ±20 s relative to gate crossing event (before: −20 to 0 s, after: 0 to +20 s). (E) Burst occurrence before gate crossing (−20 to 0 s relative to gate crossing) as a function of the number of repeated sessions. (F) Mean burst durations before gate crossing (−20 to 0 s relative to gate crossing) as a function of repeated sessions. The 1st to 4th sessions were tested on one day, and the 5th to 8th sessions were tested on the next day. * *p* < .05, ** *p* < .01. Wilcoxon signed rank test for medians of sessions 1 and 8 (α = .05, uncorrected).

### β*_high_* modulates fast gamma activity

Lastly, we asked the role of this goal-related activity of *β*_high_ in information flow within the mPFC-BLA. Prefrontal *β*_high_ has been shown to inhibit local spiking activities (Miller et al., 2018) and direct the flow of top-down information while interrupting task-irrelevant feedforward sensory input (Bastos et al., 2020b). To test whether *β*_high_ has an inhibitory effect on the local network, we analyzed fast gamma activities (60–120 Hz, γ*_fast_*) as a proxy for local excitability in the cortex (Ray and Maunsell, 2011). Overall, the amplitudes of *θ*_high_, *β*_high_ and γ*_fast_* increased during the *flight* compared to the baseline (**Figure 8A**). It is noteworthy that the augmented level of γ*_fast_* was significantly smaller in the gate-toward *flight* compared to the corner-toward *flight* in BLA (*p* = .094 in the mPFC; *p* = .002 in the BLA), whereas the augmented level of *β*_high_ was significantly higher in gate-directed *flight* in mPFC and BLA (*p* = .004 in the mPFC; *p* = .000 in the BLA) (**Figure 8A**).

**Figure 8.**
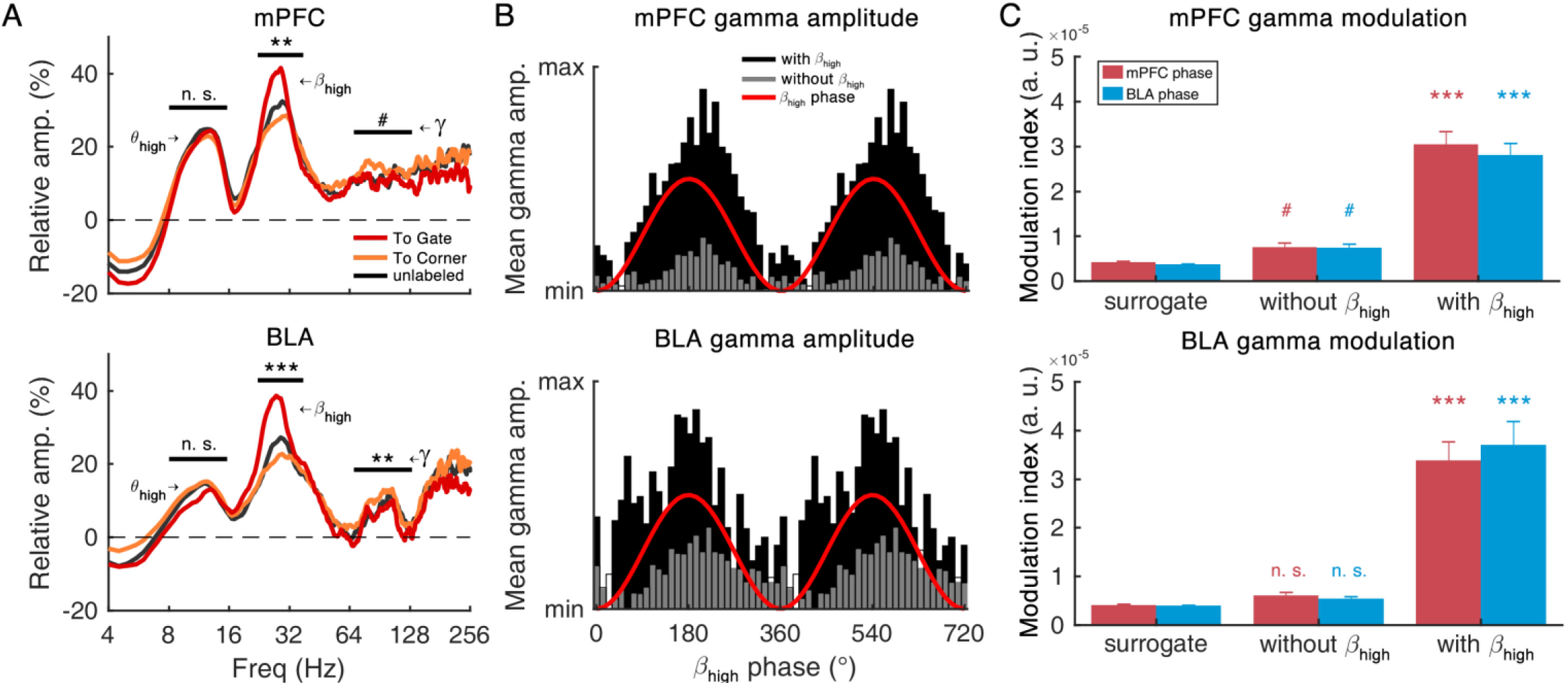
High beta waves modulate cortical excitability. (A) Amplitude spectra of flight bouts labeled by the endpoint of trajectory. Statistical tests were performed against the null hypothesis that the two medians (To Gate vs. To Corner) are equal (*post hoc* comparison after Kruskal– Wallis test). (B) Example beta-gamma phase-amplitude coupling from one mouse. Red lines denote the phase of high beta rhythm. (C) Comparison of phase-amplitude coupling in the absence of high beta waves (i.e., without) and the presence of high beta waves (i.e., with). # *p* < .10, * *p* < .05, ** *p* < .01, *** *p* < .001.

To investigate an interaction between *β*_high_ and γ_fast_, we analyzed the cross-frequency coupling between the *β*_high_ phase and the γ_fast_ amplitude. Rather than the haphazard inhibition of high-frequency activity, the γ_fast_ amplitude was temporally modulated by the *β*_high_ phase and this modulation (i.e., phasic inhibition) was higher in the presence *β*_high_ bursts (**Figure 8B**). The amplitude of fast gamma was the strongest around the peak of the *β*_high_ phase compared to that near the trough. Such phasic inhibition of fast gamma by *β*_high_ was not significant during the absence of *β*_high_ (**Figure 8C**), measured by the phase-amplitude coupling modulation index (Tort et al., 2009) both for mPFC gamma (*p*s > .0612 without *β*_high_; *p*s < .001 with *β*_high_; *post hoc* comparison with Kruskal–Wallis ANOVA, *p* < .001) and BLA gamma (*p*s > .2651 without *β*_high_; *p*s < .001 with *β*_high_; *post hoc* comparison with Kruskal–Wallis ANOVA, *p* < .001). These results suggested that the role of *β*_high_ that emerged during gate-directed *flight* was tightly linked to cortical inhibition for better processing of top-down information.

## Discussion

The primary aim of the current study was to identify the involvement of transient neural dynamics in the mPFC-BLA in supporting diverse behaviors under a naturalistic threat. We identified a *freezing*-related rhythm (*θ*_low_) and *flight*-related rhythms (*θ*_high_ and *β*_high_) in the mPFC and BLA. Just as freezing and flight do not co-exist, these rhythms were mutually exclusive. Within the network, mPFC led BLA in *θ*_low_ and *β*_high_ bands, whereas BLA led mPFC in *θ*_high_ band. Unlike theta rhythms, *β*_high_ emerging during safe zone-oriented escaping behaviors presented several properties implicating its relevance to top-down processing. Our findings on the mPFC-BLA oscillatory network during freeze-or-flight behaviors shed light on the unnoticed aspect of the process that helps flexible behavioral switching.

### Low theta in mPFC-BLA during freezing behavior

It is noticeable that our experimental setting does not account for fear learning. So far, *θ*_low_ rhythm in mPFC-BLA during freezing behavior has been observed in the Pavlovian fear conditioning paradigm (Bagur et al., 2021; Davis et al., 2017; Karalis et al., 2016; Moberly et al., 2018; Ozawa et al., 2020). For this reason, mPFC-BLA *θ*_low_ has often been interpreted as an index of successful fear memory with evidence that the duration of freezing, a primary index of memory performance, was positively correlated with *θ*_low_ power (Karalis et al., 2016). Furthermore, optogenetic interrogation showed that freezing could be induced or blocked by in-phase or out-of-phase 4 Hz stimulation (Karalis et al., 2016; Ozawa et al., 2020). However, it would be more compelling to relate *θ*_low_ with an expression of fear through freezing rather than memory processes. In our data, the freezing duration did not change as the session repeated indicating no memory effect here (**Figure S2A**), but the positive correlation between freezing duration and *θ*_low_ power was observed as well (**Figure S4C**). Likewise, *θ*_low_ emerged significantly in advance of freezing.

In this study, the question was whether *θ*_low_ is a necessary condition for freezing or not. Our event occurrence analysis showed that only a half of freezing was accompanied by *θ*_low_ bursts (**Figure 3E**), revealing that *θ*_low_ is not a necessary condition for freezing. Although LFP is a limited measure in reflecting all collective firings of neuronal ensembles, it is noteworthy that the temporal overlap ratio is low (∼30% only in the *θ*_low_-accompanied freezing. We speculate that the physiology associated with freezing may generate *θ*_low_ rhythm in the brain. Recent studies have shown that prefrontal *θ*_low_ during freezing is driven by respiration: during fear-conditioning freezing, mice breath regularly at ∼4 Hz, modulating PFC *θ*_low_ rhythm (Moberly et al., 2018) and furthermore, *θ*_low_ rhythm entrained by respiration regulates the freezing behavior (Bagur et al., 2021). Still, it is unlikely that only respiration drives *θ*_low_ rhythm because breathing never stop. Previously, an absence of *θ*_low_ rhythm was observed during non-freezing periods (Bagur et al., 2021; Moberly et al., 2018). A disruption of the olfactory bulb was reported to impair the respiration-related *θ*_low_ rhythm (Bagur et al., 2021; Karalis and Sirota, 2022; Kum et al., 2019; Moberly et al., 2018), but did not abolish freezing behavior (Bagur et al., 2021; Karalis and Sirota, 2022). Therefore, it would be appropriate to regard freezing and *θ*_low_ as parasympathetic responses (e.g., accompanying physiological phenomena to induce behavioral inhibition) to acute threats, which mutually interact with each other.

Nevertheless, *θ*_low_ participates in neural processes that promote freezing behavior. For instance, the in-phase stimulation of mPFC and BLA in a fear-conditioned context recruited functionally distinct ensembles, some of which are associated with memory retrieval for freezing (Ozawa et al., 2020). Then, the question is what cognitive functions *θ*_low_ instigate. Given that the primary role of slow oscillations like *θ*_low_ has been thought to facilitate long-range neural communication across distant cortical areas (Von Stein and Sarnthein, 2000), the mPFC-BLA *θ*_low_ may reflect the recruitment of cortical ensembles that are needed for successful execution of freezing behavior. First of all, considering that freezing behavior promotes the perception of coarse visual scenes (Lojowska et al., 2015) in humans, the recruitment of *θ*_low_ during freezing behavior may be linked to the internal attention for upcoming the threat cue. Previous findings support this idea, which suggests the PFC of mice forms a long-range network in 4-7 Hz with sensory areas such as the visual cortex (Han et al., 2019) and the olfactory bulb (Bagur et al., 2021; Moberly et al., 2018). Secondly, as reviewed by Roelfs (Roelofs, 2017), freezing requires an active preparation process for upcoming action rather than a passive response to the threat. To that end, the brain implements a parasympathetic brake on the motor system during freezing. Accordingly, Fujisawa & Buzsáki (Fujisawa and Buzsáki, 2011) reported that rat mPFC forms 4-Hz rhythmic network of the ventral tegmental area, one of the major input structures of basal ganglia, to facilitate the active communication between these two structures. That is to say, the successful execution of freezing depends critically on efficient brain connection with motor and sensory systems, and *θ*_low_ may mediate it by forming a long-range coherent network. It still remains an open question whether this *θ*_low_ acts as a primary driver of the freezing behavior or simply an accompanying phenomenon, however, the *θ*_low_-band resonance of mPFC with sensory- and motor systems at least won’t be an impediment to effective neural communication.

### High theta in mPFC-BLA during flight behavior

In fear conditioning experiment, the *θ*_high_ rhythm in mPFC-BLA has been interpreted as an expression of safety or the index of fear extinction (Davis et al., 2017; Ozawa et al., 2020). However, in our naturalistic threat experiment, the mPFC-BLA *θ*_high_ emerged upon the introduction of the predator robot and decreased after the mice escaped to the safe zone. Therefore, it is reasonable to consider the mPFC-BLA *θ*_high_ rhythm as a network state facilitating the cognitive processes for appropriate reaction in an anxiogenic environment. For instance, an efficient escaping is a high-level cognitive process based on functions like perception, prediction, spatial coordination, and execution (Roelofs, 2017). Likewise, the fear extinction involves the learning and inferring of the causal relation between passively observed events. These complex cognition and behavior will arise through dynamic recruitment of large-scale brain network and the mPFC-BLA *θ*_high_ rhythm may be a signature of this process. Yet, the brain states characterized by engaging distinct functional network needs to be identified over time to link the cognitive process and behavioral changes.

In a separate, the *θ*_high_ in mPFC-BLA here resembles the functional characteristic of hippocampal theta rhythm. The rhythmicity was prominent during locomotion. It has been well-known that hippocampal theta power is positively correlated with the movement speed (Ahmed and Mehta, 2012), which was replicated in our data (**Figure 4B, 4D**). Moreover, *θ*_high_ rhythms in the mPFC and BLA behaved like having a common driver. The time delay of *θ*_high_ rhythms in two regions was infinitesimal (∼2 ms, BLA led mPFC), whereas that of *θ*_low_ rhythm was not negligible (*∼*12 ms, mPFC led BLA). This (near) zero-lag synchrony between long-range ensembles can be implemented through dynamical relaying aided by the common driver (Uhlhaas et al., 2009; Vicente et al., 2008). Furthermore, our results of flight-specific *θ*_high_ is consistent with the idea that mPFC-BLA-hippocampus (HPC) triodes interact while processing fear and safety (Dejean et al., 2015; Herry et al., 2010; Maren and Quirk, 2004), and their functional interactions depend on oscillatory long-range synchronization (Bocchio and Capogna, 2014; Stujenske et al., 2014). In sum, *θ*_high_ in mPFC-BLA in a part of the cortico-hippocampus network promotes reconfiguration of the circuits for a successful escape.

### Mutual exclusivity of *θ*_low_ and *θ*_high_ rhythms during *freezing* versus *flight*

Just as freezing and flight cannot exist simultaneously, the coexistence of freezing- (i.e., *θ*_low_) and flight-specific (i.e., *θ*_high_) oscillatory states was rarely found. This mutual exclusivity (i.e., winner-takes-all) of distinct rhythms may provide an efficient cortical implementation of flexible switching, especially for the two opposite behaviors like freezing and flight. What circuit motif could support this efficient switching between these two non-overlapping rhythms?

One answer could be the competitive interaction of distinct subpopulations that engages distinct rhythmogenic processes. In fact, the distinct neuronal populations related to fear behavior are relatively well documented in the amygdala (for review, see (Janak and Tye, 2015)): In the CeA, Fadok et al. (Fadok et al., 2017) defined two types of inhibitory neurons regulating dimorphic defensive behaviors and their competitive interaction. In particular, corticotropin-releasing factor neurons in CeA promote conditioned flight, somatostatin-positive neurons in CeA promote conditioned freezing, and the two populations competitively inhibit each other and balance the selection of appropriate defensive behavior (Fadok et al., 2017). Still, the question remains how specific populations are recruited and how others are rapidly dismissed. Our finding hints how neuronal circuits achieve it through the rhythms (and selective resonance). In order to achieve the behavior robustness, the formation of corresponding neural assemblies will occur in a mutually exclusive way, reasonably inhibiting one another to guarantee this exclusive formation, as inferred from the exclusive relationship of *θ*_low_ and *θ*_high_ observed here.

Moreover, at present, it remains unclear whether these distinct rhythms causally regulate the interactions of these subpopulations or result from them. Of note, there is a report that inhibition of parvalbumin-positive (PV+) interneurons disrupt the balance between *θ*_low_ and *θ*_high_ by increasing *θ*_low_ and decreasing *θ*_high,_ which was shown to result in longer conditioned freezing behavior (Davis et al., 2017), indicating that hub-like inhibitory neurons can regulate the oscillations. More recently, Ozawa et al. (Ozawa et al., 2020) demonstrated that optogenetic entrainment of *θ*_low_ and *θ*_high_ oscillations in the PV+ population recruited distinct phase-locking ensembles in the BLA, suggesting that these ensembles have selective resonance such that exogenously driven rhythms can determine which population participates. However, it is unlikely that a locally driven oscillation itself switches the entire behavioral state and controls the oscillations at the mesoscopic level. Rather, it has been considered that coordination of inter-areal interplay underlies these switches in behavioral states and specific projections based on the exhibited behavior playing a role in the robust recruitment of pre- and postsynaptic groups (Senn et al., 2014; Tovote et al., 2016). In line with this, Senn et al. (Senn et al., 2014) found a distinction between mPFC-projecting BLA neurons during freezing (prelimbic) versus non-freezing (infralimbic) behaviors. More recently, Amir et al. (Amir et al., 2018) reported that the distinct types of principal neurons in the BLA are entrained to local oscillations depending upon their projection sites. Therefore, we speculate that the distinct theta rhythms facilitate the robust formation of interregional neuronal assemblies via mutual inhibition of behavior-specific neuronal assemblies. Nonetheless, an exact mechanism underpinning the rapid transition is still needed.

### Prefrontal-driven beta oscillations and top-down processing

*β*_high_ rhythmicity during flight, especially during the safe-zone-directed escape behavior, exhibited prefrontal-to-amygdala directionality as well as a top-down aspect with power around the gate area that increased across trial repetitions. Beta oscillations in the cerebral cortex have been associated with motor function (Brovelli et al., 2004; Feingold et al., 2015; Pfurtscheller et al., 1996) and the maintenance of status quo (Engel and Fries, 2010); however, studies of the beta rhythm in the non-sensorimotor domain are relatively rare, especially in rodents. Only a limited number of studies have investigated the *β*_high_ in rodent mPFC and BLA; notwithstanding, the beta rhythm in the rodent mPFC-BLA circuit has been reported to be tightly linked to top-down processes. Specifically, Samson et al. (Samson et al., 2017) found transient beta-rhythmic activity in the rat BLA during reward expectation (i.e., the time before reward after lever pressing) in an operant conditioning task, resembling our finding of spatially selective recruitment of *β*_high_ around the gate area in relation to reward expectation (i.e., exiting the threat zone). Moreover, Zhang et al. (Zhang et al., 2019) reported beta rhythmicity recruited during the recognition phase in a spatial working memory task, showing its putative role in top-down processing in the mouse mPFC for successful goal-directed behavior. Converging evidence has indicated the pivotal role of amygdala-prefrontal interactions in valence coding and the emotional learning (Pignatelli and Beyeler, 2019; Yizhar and Klavir, 2018); however, exact circuit mechanisms in this complex network are far from our understanding. Future studies may address how the brain does this by dissecting the computational significance of oscillations transiently emerging during top-down processes.

Numerous findings from human and nonhuman primate brains have suggested that prefrontal-driven beta rhythms play a key role in top-down cognitive processes, such as working memory (Engel and Fries, 2010; Lundqvist et al., 2018). Recent theoretical work on the cortical beta rhythm has proposed that it contributes to the *prediction* of sensory information by inhibiting fast feedforward neural activities (i.e., gamma and spiking activity) for better processing of information that matches the expected input (Bastos et al., 2020a). Although our recording of LFPs did not allow investigation of the laminar specificity that the original literature emphasized (Bastos et al., 2012; Miller et al., 2018), the properties of beta rhythms match well with the concept of predictive routing in our findings based on (1) increases with trial repetition, (2) prefrontal-leading directionality of information flow, and (3) silencing effects over local excitability (i.e., cross-frequency interaction between beta and fast gamma). In this interpretation, the role of the beta rhythm during the defensive behavior is somewhat like ‘priming’ safety-related information in relevant brain regions to allow easier processing of an upcoming predicted event (i.e., gate opening), while silencing other escape-irrelevant sensory information (i.e., distractors) held in fast-frequency feedforward activities. Desynchronization of the beta rhythm after crossing the gate also supports this concept, as the prediction is no longer needed in the safe zone. A remaining question is the trigger of these prefrontal-driven oscillations, which could provide a clue to the spontaneous generation of goal-directed behaviors.

### Oscillatory gear-shifting supports flexible switching of different behaviors

Here, we report distinct network states of *θ*_low_, *θ*_high_, and *β*_high_ in mPFC-BLA circuit dynamically switches upon threat context and defensive behaviors. Considering the complex anatomical connections that both the mPFC and BLA possess, such switching in oscillatory frequencies may provide a cost-effective way to form a flexible network in a timely manner with other resonant brain areas, minimally affecting the anatomical architecture of the whole system. In this selection of one over multiple frequency alternatives, the brain may coordinate functional ensembles which are resonantly distributed over the brain cost-effectively, and support the flexible switching among different behaviors. The computational advantage of having multiple frequency channels in neuronal circuitry has been suggested by the mechanism of *oscillatory multiplexing* (Akam and Kullmann, 2014) for selective inter-areal communication, and experimental evidence supports it, such as the distinction of hippocampal slow and fast gamma activity (Colgin, 2015; Colgin et al., 2009). However, having multiple communication routes established by distinct frequencies (i.e., oscillatory multiplexing) does not always lead to flexible switching of the network state that eventually guides behavioral flexibility. Our findings on the role of distinct oscillations in the mPFC-BLA network resemble oscillatory multiplexing such that they can be understood as a specific form of it (i.e., with a constraint for interdependence across distinct frequency-specific populations). From this point of view, the existence of nonoverlapping rhythmic states and the ability to switch among them is similar to a gear-shifting of an automobile, where one appropriate driving mode (i.e., appropriate type of defensive behavior) is chosen over the others, which transiently interconnects functionally relevant parts (i.e., different brain regions).

In sum, the brain is a complex dynamical system composed of modular and hierarchically organized circuits with various functions (Meunier et al., 2010). To deal with an acute threat, flexible shifting between freezing and flight is needed, and the related brain circuits should engage and disengage accordingly. How the brain successfully and flexibly switches between opposite behaviors has remained as an open question. Our finding of freezing- and flight-specific rhythms in the mPFC-BLA may reflect transient formations of brain networks communicating through the coherence (Fries, 2015), and the information routing of input signals which is determined by the target regimes during the state selection (Palmigiano et al., 2017). We consider these behavior-specific reconfigurations of the defensive circuit to represent transiently formed circuits for the corresponding action that steer information transfer via oscillatory synchronization with functionally relevant brain regions. In such a way, the behavior of freeze-or-flight can be quickly implemented to coordinate the cognitive systems orchestrating relevant circuits for perception and action given ongoing task demands.

## Acknowledgement

We thank the members of the Computational Cognitive & Systems Laboratory for discussion and comments on the manuscript. This research was supported by Basic Science Research Program through the National Research Foundation of Korea (NRF) funded by the Ministry of Education (2022R1A6A3A01085957) and by grants from a National Research Foundation of Korea grant (NRF-2022R1A2C3003901) and by KIST Intramural grant (2E31511).

## Author contributions

J.H.C. supervised the study. H.-B.H. designed the concept of study, performed main data analyses and statistical tests. J.K. conducted experiments and preprocessing of the data. J.H.C. and H.-B.H. wrote the initial draft of the manuscript. H.-B.H., H.-S.S., Y.J. and J.H.C. discussed the results and interpretation of the data.

## Inclusion and diversity

One or more of the authors of this paper self-identifies as an underrepresented ethnic minority in science. One or more of the authors of this paper self-identifies as a gender minority in their field of research. We support inclusive, diverse, and equitable conduct of research.

## STAR★METHODS

KEY RESOURCE TABLE

KEY RESOURCE TABLE
| REAGENT or<br>RESOURCE | SOURCE | IDENTIFIER |
| --- | --- | --- |
| **Experimental Models: Organisms/strains** |  |  |
| **B6.Cg-Tg(Thy1-COP4/EYFP)18Gfng/J Mice** | The Jackson laboratory | JAX: 007612 |
| **Software and Algorithms** |  |  |
| **U-Net** | Ronneberger et al., 2015<br>(Ronneberger et al., 2015) | DOI: 10.1007/978-3-319-24574-4_28 |
| **Keras** | https://keras.io/ | Keras 2.2.4 (backend: TensorFlow 1.13.1) |
| **MATLAB** | The MathWorks Inc. | Version 2019 |
| **Python** | https://www.python.org/downloads/ | Python 3.7.3 |
| **StreamPix** | Norpix | StreamPix 7 |
| **Other** |  |  |
| **Tungsten microelectrode** | A-M Systems | #796000, #575400 |
| **CBRAIN** | DOI: 10.1126/sciadv.abb9841 | Available upon request |
| **Spider robot** | Academy Plastic Model Company | #18143A |
| **Fiber optic cannula** | Hangzhou Newdoon Technology | FOC-C-1.25-200-7-0.37 |
| **CMOS camera** | Teledyne Lumenera | Lt225 |

## RESOURCE AVAILABILITY

### Lead contact

Further information and requests for reagents and resources may be directed to and will be fulfilled by the Lead Contact, Dr. Jee Hyun Choi.

### Materials availability

This study did not generate new unique reagents or mouse lines.

### Data and code availability

Data reported in this paper will be shared by the lead contact upon request. Any additional information required to reanalyze the data reported in this paper is available from the lead contact upon request.

## EXPERIMENTAL MODEL AND SUBJECT DETAILS

We used the data collected from our previous study (Kim et al., 2020) including LFP data from mPFC which was not analyzed in the original publication. Our previous study was performed after approval by the Korea Institute of Science and Technology (KIST) Animal Care and Use Committee (permit number: 2019-095) and under compliance with the National Institute of Health Guidelines for minimizing the pain and discomfort of animals. A total of eight, healthy, 8-week-old, male B6 mice (Thy1-COP4-EYFP) weighing more than 25 g were used for electrode-implantation surgery, and experimental procedures were performed 2-3 weeks after surgery. Mice were kept individually at the KIST Animal Facility on a 12-h light/dark cycle with *ad libitum* access to food and water.

## METHOD DETAILS

### Surgery

To obtain *in vivo* electrophysiological data from mPFC and BLA in freely-behaving animal, all mice underwent surgery for chronic electrode implantation. A ketamine-xylazine cocktail (120 and 6 mg/kg, respectively) was used to induce general anesthesia during surgery. To assess the mPFC, a single strand of PFA-coated tungsten electrode (flat tip, 76.2-µm inner diameter, Cat# 796000, A-M Systems, WA, US) was implanted in the prelimbic subregion of the right mPFC (anterior-posterior: 1.4 to 1.8 mm; medial-lateral: 0.3 to 0.5 mm; dorsal-ventral: -2.2 to -2.5 mm). To evaluate the amygdala, an epoxy-insulated tungsten electrode (8 deg tapered tip, 125-µm inner diameter, Cat# 575400, A-M Systems, WA, US) coupled with fiber optic cannula (FOC-C-C-1.25-200-7-0.37, Newdoon, China) for intraoperative stimulation procedure that targeted the basolateral nucleus (anterior-posterior: -1.9 to -1.6 mm; medial-lateral: 2.5 to 3.1 mm; dorsal-ventral: -4.2 to -4.9 mm) was implanted. An intraoperative stimulation protocol was performed to ensure successful targeting of the BLA electrode implantation as previously described (Kim et al., 2020). Two screw electrodes for the ground and reference were implanted onto the occipital bone, and all electrodes were fixed and secured with dental cement (Vertex Self-Curing, Vertex Dental, Zeist, The Netherlands). Experimental procedures were performed after at least 2 weeks of recovery period.

### Threat-and-escape paradigm

All experiments were conducted in a square-shaped, custom-built arena divided diagonally. The arena (60 cm × 60 cm) was divided into two zones (safe and threat zones), separated by a diagonal wall with a retractable gate. The arena was located in a sound-proof and electrically shielded room under dim light (room temperature: 22°-24°). After thirty to ninety minutes of habituation (mouse freely explored the safe and threat zones), a consecutive four-stage experimental protocol began: (Stage 1) the animal freely explored the threat zone without a threat present for 60 s; (Stage 2) a spider robot (Model 18143A, Academy Plastic Model Co. Ltd., South Korea) was placed into the arena and chased the mice for 60 s; (Stage 3) the gate to the safe zone was opened so that the animal could escape from the robot (60 s); (Stage 4) the robot was removed from the arena so that the animal could freely explore both the safe and threat zones (60 s). Sessions of the 4-minute protocol were repeated 8 times (4 sessions per day, two consecutive days). All mice were implanted with LFP electrodes and carried CBRAIN headstage on its own head.

### Position tracking of the mice and spider robot

The position tracking of mouse body was performed using a CNN of U-Net architecture (Ronneberger et al., 2015) by estimating the center of mass of the body area. First, the frames of the video file (1000 × 1000 pixels, 30 fps) were extracted and converted to low-resolution gray frames (448 × 448 pixels). Second, 500 frames were randomly sampled to produce a training set, and the corresponding 500 binary masks (mouse body area coded as 1, otherwise 0) were manually generated. Third, the U-Net was implemented through Keras 2.2.4 in the Python 3 environment (dropout = 0.2, number of filters = 8, ReLU activation, number of output class = 1, output activation = sigmoid, number of convolution/deconvolution blocks = 4). This CNN takes an image matrix (448 x 448 pixels) as input to produce a probability map (448 x 448 pixels) as output. Then, the output probability map was converted into a binary mask (threshold: probability of 0.4). Finally, the mouse position was determined by calculating the center of mass in the binary mask and rescaled to physical units in *cm*. Position tracking for the spider robot was performed in the same way, with a smaller number of training sets (100 frames).

### Quantification and statistical analysis

#### Behavior state labeling

A total of four behavior states (*flight*, *freeze*, *explore,* and *groom*) were identified using semiautomatic image processing. To label *freeze*, we measured the amount of bodily motion by counting the proportion of changed pixels in the time derivative of the mouse body mask frame. A frame series of slow motion (thresholding at lower 30%) was identified, and only bouts longer than 200 msec were included. To label *flight*, locomotion speed obtained from body mask centroids was used. First, we detected positive peaks in the smoothed speed vector (smoothing kernel size = 500 msec, minimum peak value = mean plus 1.96 standard deviation). Then, two negative peaks were detected: one from the left side (i.e., past) of the positive peak for *flight* initiation and the other from the right side (i.e., future) of the positive peak for *flight* termination. After automatic labeling, visual screening was performed to exclude non-defensive behaviors that have similar features as *freeze* (e.g., a sleep-like state with slow head nodding) or *flight* (e.g., locomotion toward the robot for investigation). To label *explore*, a similar method was used but with a less strict threshold (minimum peak value = mean plus 1.28 standard deviations). The behavior of *groom* (e.g., self-soothing, fur cleaning, etc.) was manually identified by the experimenter. Sequential events with the same behavioral states with marginal intervals (shorter than 200 msec) were regarded as one event and merged. All labeled events were visually confirmed by the experimenter.

#### LFP data analyses

LFP signals were obtained with CBRAIN wireless headstage (2.6 g in total, INTAN RHD 2216 amplifier board inside) then transmitted to recording PC through Bluetooth interface (nRF52832, Nordic Semiconductor), and saved at 1024 Hz with custom-written MATLAB (R2019a, Mathworks, MA, US) acquisition software. The LFP signals from the mPFC and BLA were preprocessed (i.e., highpass filtered at 0.1 Hz, bandstop notch filtered at 60 Hz, and artifacts removed) before spectral analyses.

##### Spectral analysis

Amplitude and cross-spectral density of LFPs were estimated using fast Fourier transform (FFT) with MATLAB built-in *fft.m* function with a 2048-size sliding (0.1-s step) Hanning window to draw a time-frequency representation of the LFP signal.

##### Instantaneous amplitude

The instantaneous amplitudes of LFPs were calculated via the Hilbert transform (Pikovsky et al., 2002) using the *hilbert.m* function of MATLAB after bandpass filtering (3–7 Hz for *θ*_low_; 8–14 Hz for *θ*_high_; 22–34 Hz for *β*_high_).

##### Cross-frequency coexistence and exclusion matrix

A cross-frequency coexistence matrix was calculated from a spectrogram for each frequency bin (threshold: upper 5% for coexistence, lower 5% for exclusion) (see Figure 1b in (Han et al., 2017)) by quantizing it into binary time series. Taking a pair of seed (*f*_1_) and target (*f*_2_) frequencies from binary time-frequency spectrogram(s), the conditional probability *p*(∃*f*_2_|∃*f*_1_) was calculated and assigned as the cross-frequency coexistence matrix. Likewise, the cross-frequency exclusion matrix was calculated with *p*(∄*f*_2_|∃*f*_1_). The maps were produced by averaging the matrices. We determined significant increases or decreases in the value by calculating the average of the conditional probabilities with a temporally shuffled seed array and assigning it as the chance level. The detailed calculation procedure is summarized in **Figure S5**.

##### Cross-correlation analysis

To determine the lead-lag relationship of LFPs between two regions, the cross-correlation function was calculated. First, LFP signals were bandpass filtered using a 5th-order Butterworth filter. The frequency bin was set to 1–150 Hz with a 1-Hz interval, and the bandwidth was set to a quarter of the center frequency (i.e., ƒ±ƒ/8 Hz). Second, a cross-correlation function was obtained using the MATLAB built-in *xcorr.m* function (considered max lag size = 3 cycles) with the same application of a moving window (2048 size, 0.1-s step). Finally, a time lag with maximal (correlation) or minimum (anti-correlation) value, τ, was taken from the cross-correlation function. After taking τ, its unit was converted into radians for comparison across frequencies.

##### Phase-amplitude coupling analyses

To analyze cross-frequency phase-amplitude coupling (PAC), raw LFP signals were Hilbert-transformed after bandpass filtering (phase frequency bin = 2 to 64 Hz with a 2-Hz interval (1-Hz bandwidth); amplitude frequency bin = 16–32 Hz with a 2-Hz interval (2-Hz bandwidth), 32–64 Hz with a 4-Hz interval (4-Hz bandwidth), 64–128 Hz with a 16-Hz interval (8-Hz bandwidth), 128–256 Hz with a 32-Hz interval (32 Hz-bandwidth), and 256–512 Hz with a 64-Hz interval (64-Hz bandwidth)). We used Tort et al.’s method to calculate the modulation index (Tort et al., 2009) with a phase bin size of 30.

##### Spatial distribution of LFP amplitude

The position of the mice was binned into a two-dimensional square grid (112 x 112 pixels), and the amplitude extracted from the spectrogram in each position bin was averaged to obtain the spatial distribution of the frequency components. For baseline correction, the amplitude in each position bin was divided by the session means across all stages and converted into a percentage scale. Averaging kernels were defined as sector-shape masks (radius = 30 pixels) for the corner area and a rectangle-shape (*width*:*height* = 15:24 pixels) mask rotated along a diagonal axis (45°) for the gate area.

##### Unsupervised clustering of flight trajectory

*Flight* trajectories were clustered using the *k*-means algorithm implemented in MATLAB Statistics and Machine Learning Toolbox (v11.5). For better clustering performance, achieved by avoiding large input dimensionality, the raw binary trajectory image (56 x 56 pixels, contains position information) and its time derivative (56 x 56 pixels, contains direction information) were compressed into a low-dimensional dense layer (8 nodes) using a variational autoencoder. The appropriate number of clusters (*k* = 10) was determined by a silhouette score-based goodness-of-fit (squared Euclidean distance measured by *silhouette.m* in MATLAB) reaches its maximum (silhouette score = 0.4) from a grid search (from *k* = 2 to *k* = 128, **Figure S8**).

#### Statistical testing

For statistical testing, nonparametric versions of t test (Wilcoxon signed rank test, rank sum test) and ANOVA (Kruskal-Wallis test) were used. For correlation coefficients, both Pearson’s and Spearman’s method were both used depending on the type of data analysis. Alpha level was set to 0.05.

#### Data and software availability

The datasets and the codes are available upon reasonable request.

#### Resource availability

Further information and requests for the device and resources may be directed to and will be fulfilled by the Lead Contact, Dr. Jee Hyun Choi.

## Supplementary Information

**Figure S1.**
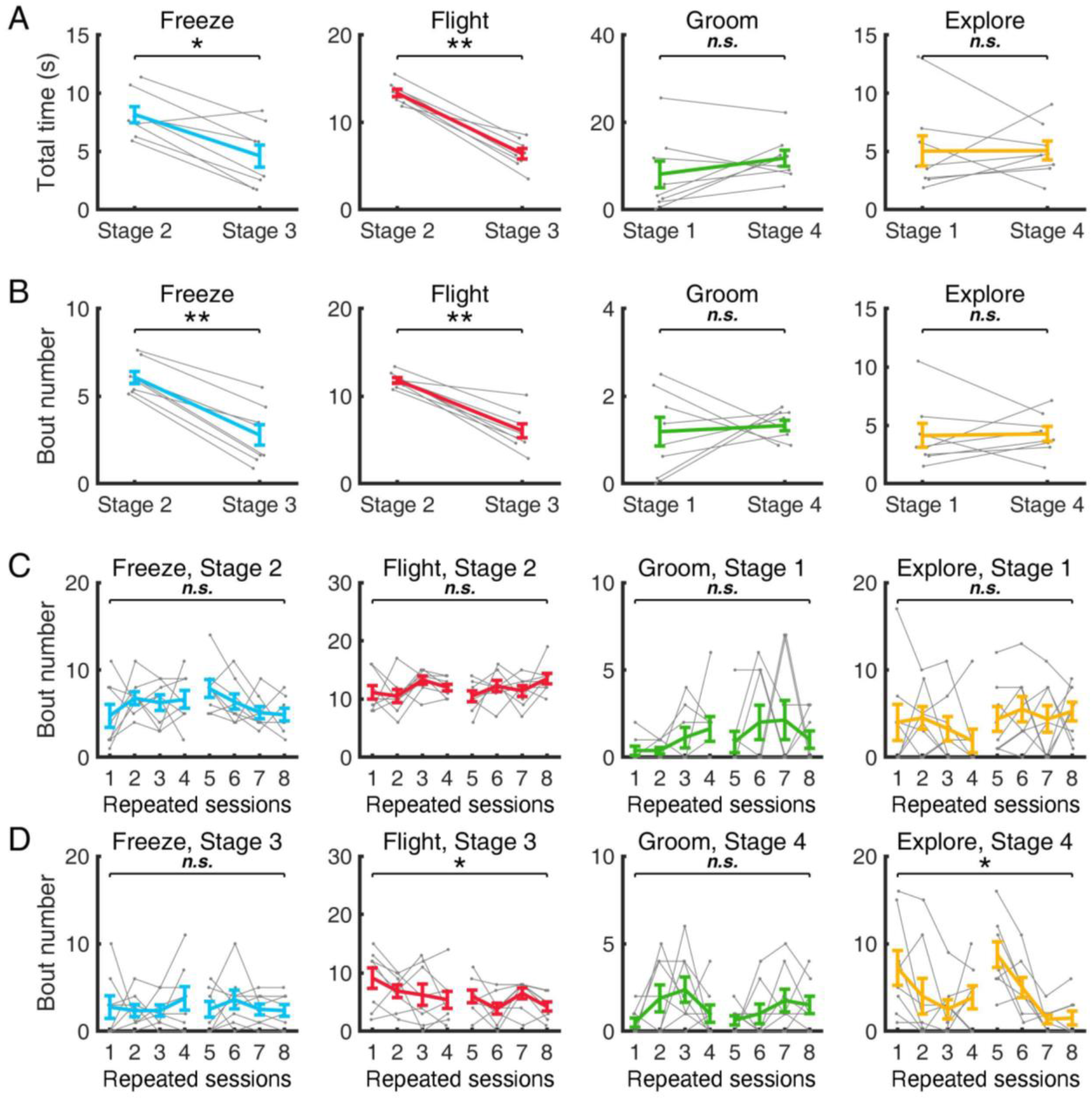
Comparison of behavioral state occurrence rates as a function of stage and across repeated sessions, Related to Figure 1. (A) Total time in each behavioral state in different stages showing that more *freezing* and *flight* bouts occurred during Stage 2 (robot attack, safe zone unavailable) than during Stage 3 (gate opened, safe zone available). The difference in total time for grooming and exploring behaviors between Stage 1 (baseline, before robot attack) and Stage 4 (no threat, after robot attack) was not statistically significant. (B) Same as (A) but for bout numbers. (C) Total bout numbers as a function of repeated sessions. There was no statistically significant difference between the first and the last session for Stage 2 (freeze and *flight*) or Stage 1 (groom and explore). (D) Same as (C) but for Stages 3 and 4, showing attenuated levels of locomotion for *flight* and exploration in the later sessions. The 1st to 4th sessions were tested on one day, and the 5th to 8th sessions were tested on the next day. * *p* < .05, ** *p* < .01, Wilcoxon signed rank test for medians of two stages (A-B) or sessions 1 and 8 (C-D) (α = .05). Gray dots denote the data from individual mouse.

**Figure S2.**
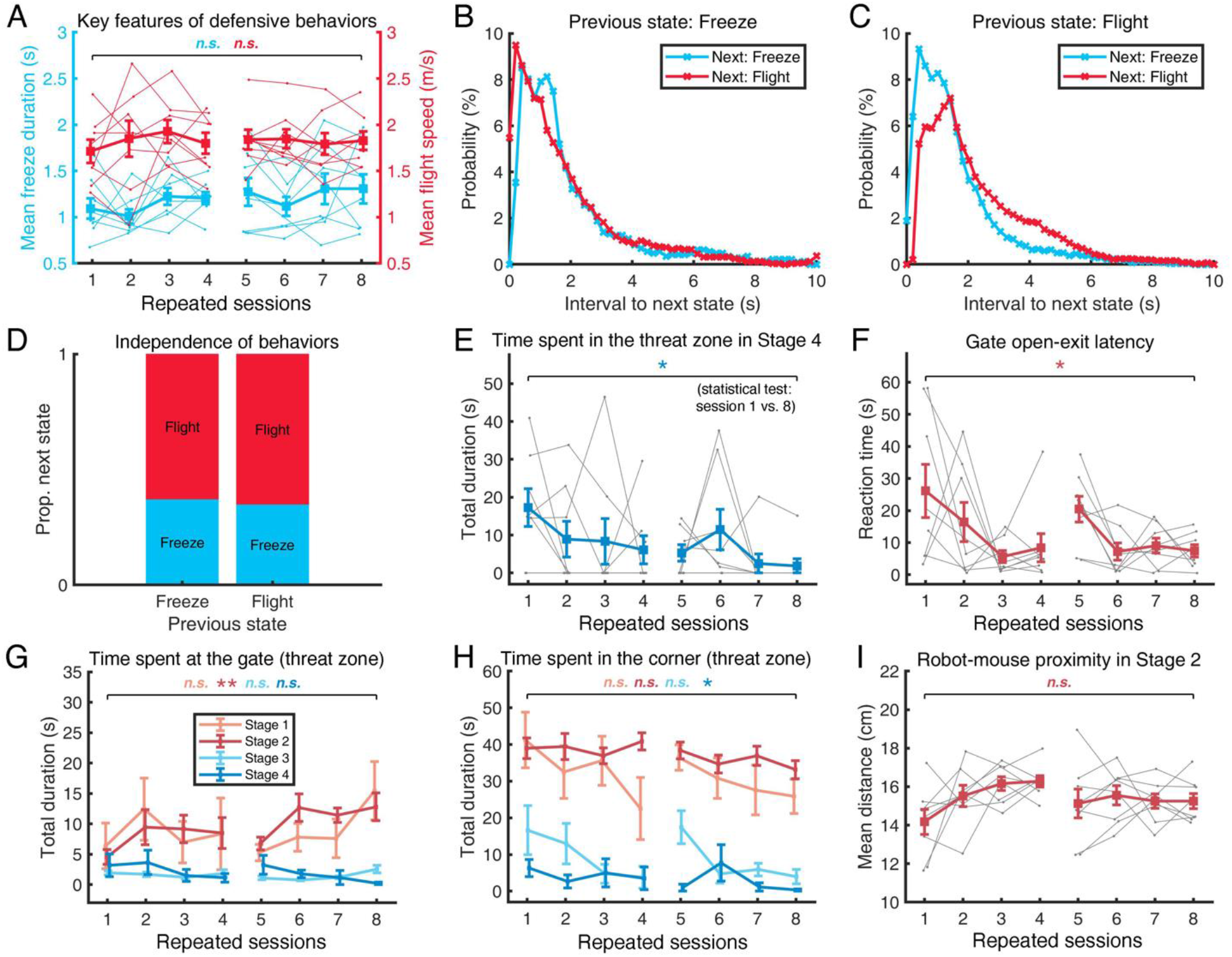
Changes across sessions in behavioral variables and state transitions during the attack period, Related to Figure 1. (A) Null effect of session repetition on the key features of defensive behaviors; *p* = .2500 for *freezing* duration, *p* = .6406 for *flight* speed. (B-C) Histogram of the interstate intervals from the *freezing* state to the next state (B) and from *flight* state to the next state (C). (D) The proportion of the next defensive action for a given action. The selection of behavior type was not dependent on the previous action. (E) The animals tended to avoid the threat zone more in the final session (mean duration = 1.89 s) than in the initial session (mean duration = 17.24 s), during the time period after the removal of the predator robot in Stage 4, showing their preference for the safe zone. (F) Likewise, the decrease in reaction time to the gate opening event in Stage 3 was statistically significant (*p* = .039), showing that animals acquired task rules as well as a preference for the safe zone. (G) This tendency to avoid the threat zone can also be found during the attack period (Stage 2), where the time spent around the gate area was significantly higher in the final session (12.81 s) than in the initial session (4.56 s), *p* = .0078. (H) The preference for the corner area during the attack period did not change (*p* = .1875). (I) This tendency for a preference for the safe zone did not originate from the change in threat level, as measured by the proximity of predator (*p* = .4609). The 1st to 4th sessions were tested on one day, and the 5th to 8th sessions were tested on the next day. * *p* < .05, ** *p* < .01. Wilcoxon signed rank test for medians of sessions 1 and 8 (α = .05).

**Figure S3.**
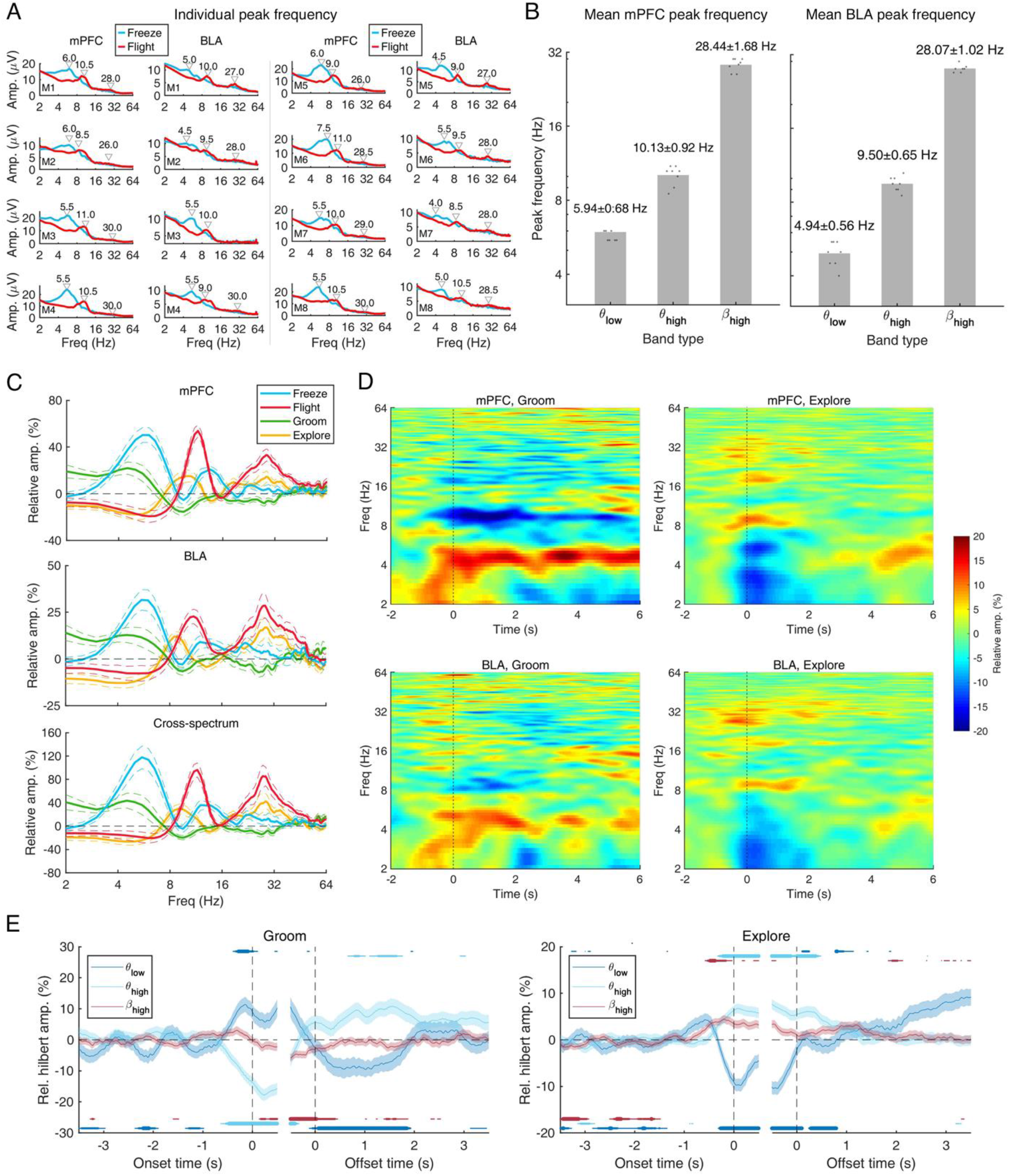
EEG amplitude spectra in mPFC and BLA, Related to Figure 2. (A) Individual peak frequency in the raw power spectrum for *θ*_low_ (3-8 Hz) during *freezing* and *θ*_high_ (8-14 Hz) and *β*_high_ (22-34 Hz) during *flight*, in the mPFC (left) and BLA (right). Peaks were detected by finding the local minima in each frequency band. (B) Mean peak frequency in the mPFC (left) and BLA (right) detected in (A). Values denote the mean and one standard deviation of the peak frequency in each frequency band. (C) Mean relative amplitude of mPFC and BLA oscillations for each behavioral state (blue: freeze; red: *flight*; green: groom; and yellow: explore) (top: mPFC; middle: BLA; bottom: mPFC-BLA cross) normalized by mean amplitudes in the baseline period. Although grooming and exploration resemble the behavioral phenotypes of *freezing* (i.e., absence of locomotion) and *flight* (i.e., presence of locomotion), respectively, strong oscillatory activities during *freezing* and *flight* were not observed. (D) Time-frequency representations of mPFC and BLA oscillations in each behavioral state. (E) Hilbert instantaneous amplitudes of narrow bandpass-filtered LFP signals relative to the onset and offset of the behavioral states of grooming and exploration. Solid bars indicate that the statistical significance against the null hypothesis that the instantaneous amplitude is equal to zero (Wilcoxon signed rank test, uncorrected), and the thickness of the bars indicates the significance level (thick: *p* < .001; middle: *p* < .01; thin: *p* < .05). * *p* < .05, ** *p* < .01 tested by Wilcoxon signed rank test against the null hypothesis that the two medians are equal.

**Figure S4.**
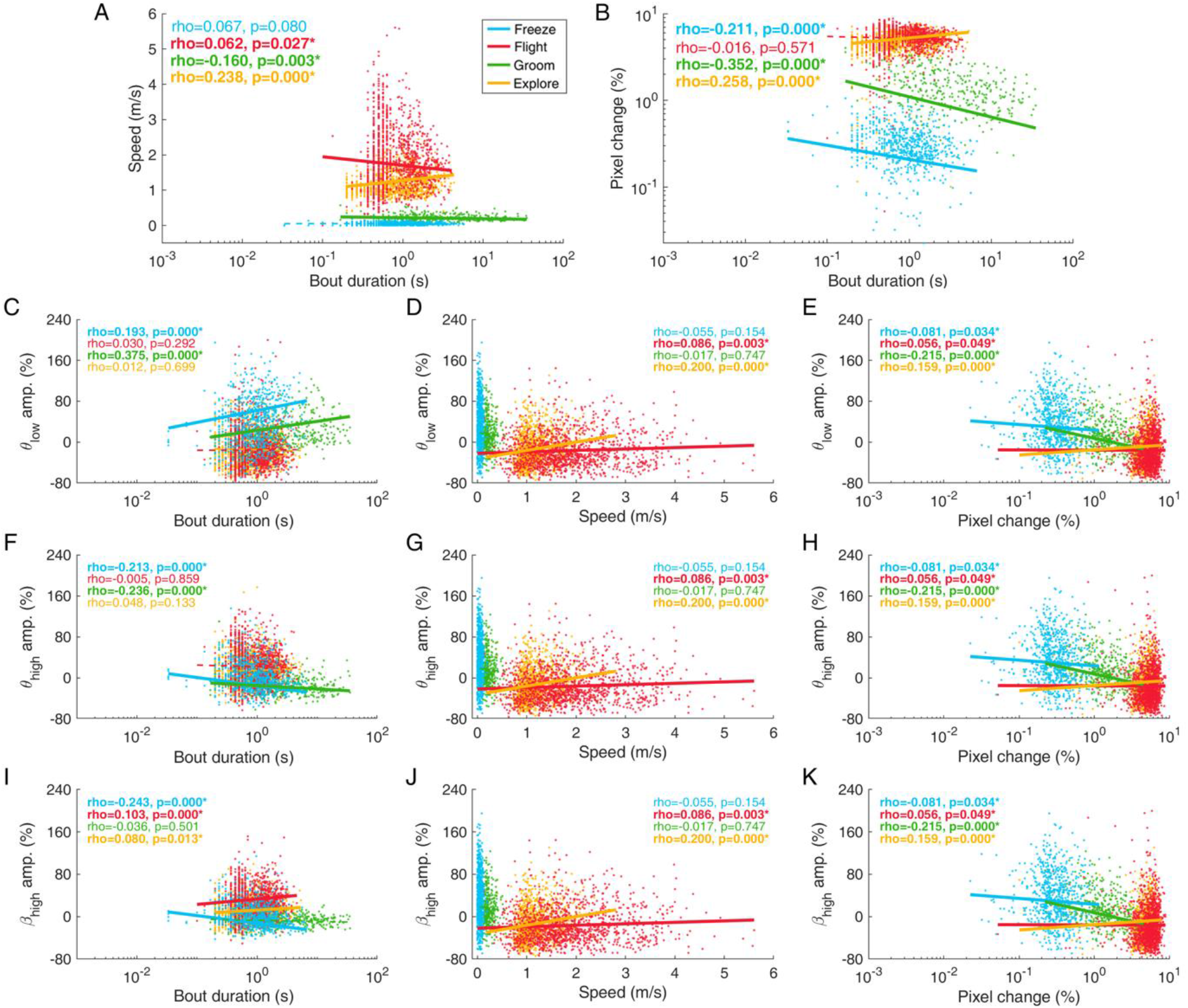
Relationships between oscillatory amplitudes in the mPFC and bout duration speed, and pixel changes, Related to Figure 2. (A-B) Scatter plots of bout duration and locomotion speed (A) and of bout duration and body pixel changes (B). (C-K) Scatter plots of behavioral variables and mPFC oscillatory amplitudes. Solid (statistically significant) and dotted lines (nonsignificant) denote the polynomial linear fitting result of x and y variables. Nonparametric correlation analysis was performed by Spearman’s rho (α = .05).

**Figure S5.**
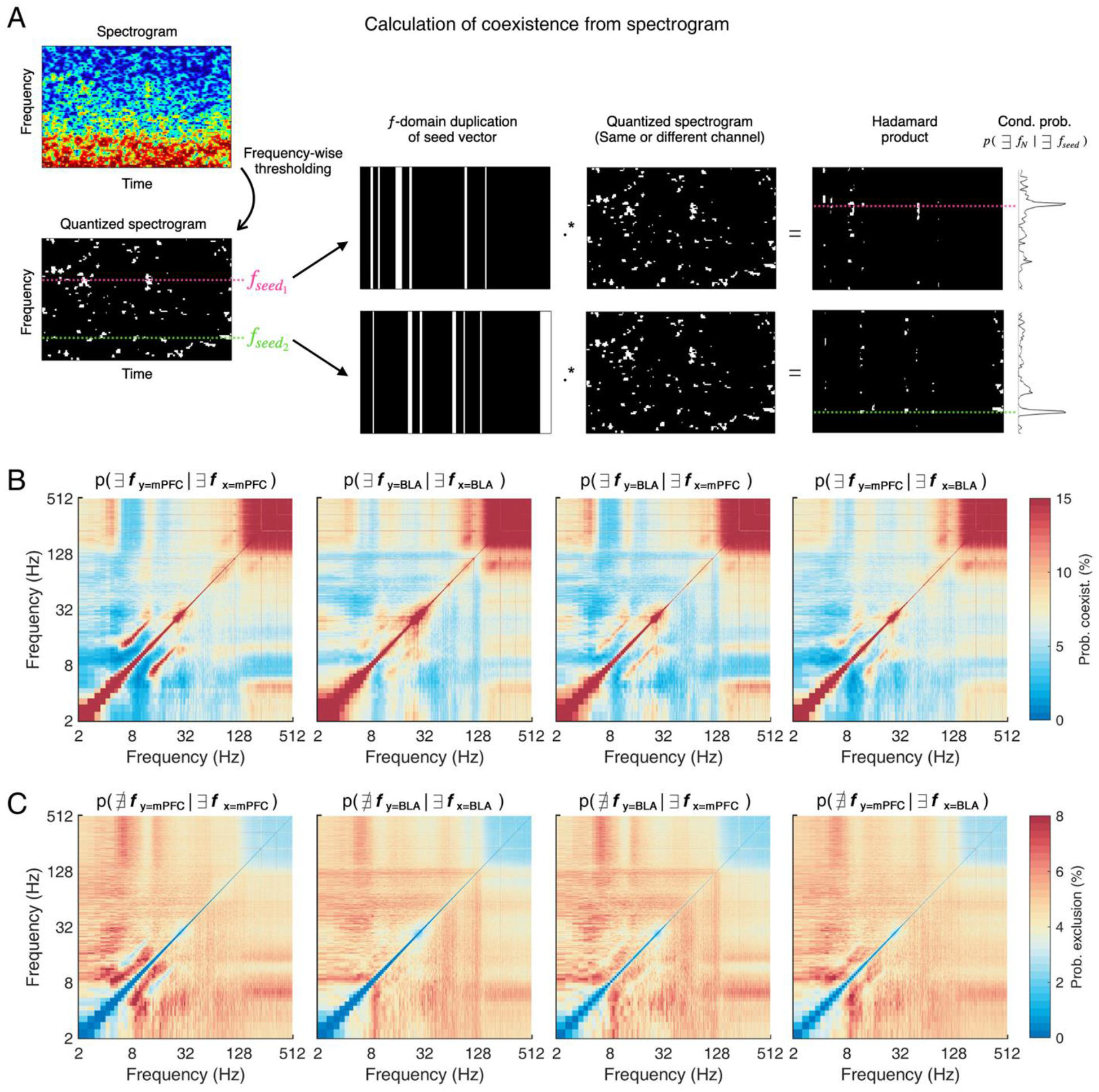
Coexistence matrix analysis in the mPFC and BLA, Related to Figure 3. (A) Schematic illustration of a quantification method for the coexistence matrix. First, the time-frequency amplitude spectrogram was quantized in a binary matrix using thresholding (threshold: upper 5% amplitude value for coexistence, lower 5% for exclusion). To calculate the conditional probability (e.g., *p*(*f_n_* = 1 |*f_seed_* = 1) for coexistence), the time series vector of seed frequency (*f_seedn_*) was selected and duplicated in the frequency domain. The Hadamard product of this duplicated vector and the original quantized spectrogram was obtained to quantify the conditional probability spectrum (rightmost black solid lines) by averaging the resultant Hadamard product in the time domain. (B) Results of the coexistence matrix analysis in the mPFC and BLA. (C) Results of exclusion matrix analysis in the mPFC and BLA.

**Figure S6.**
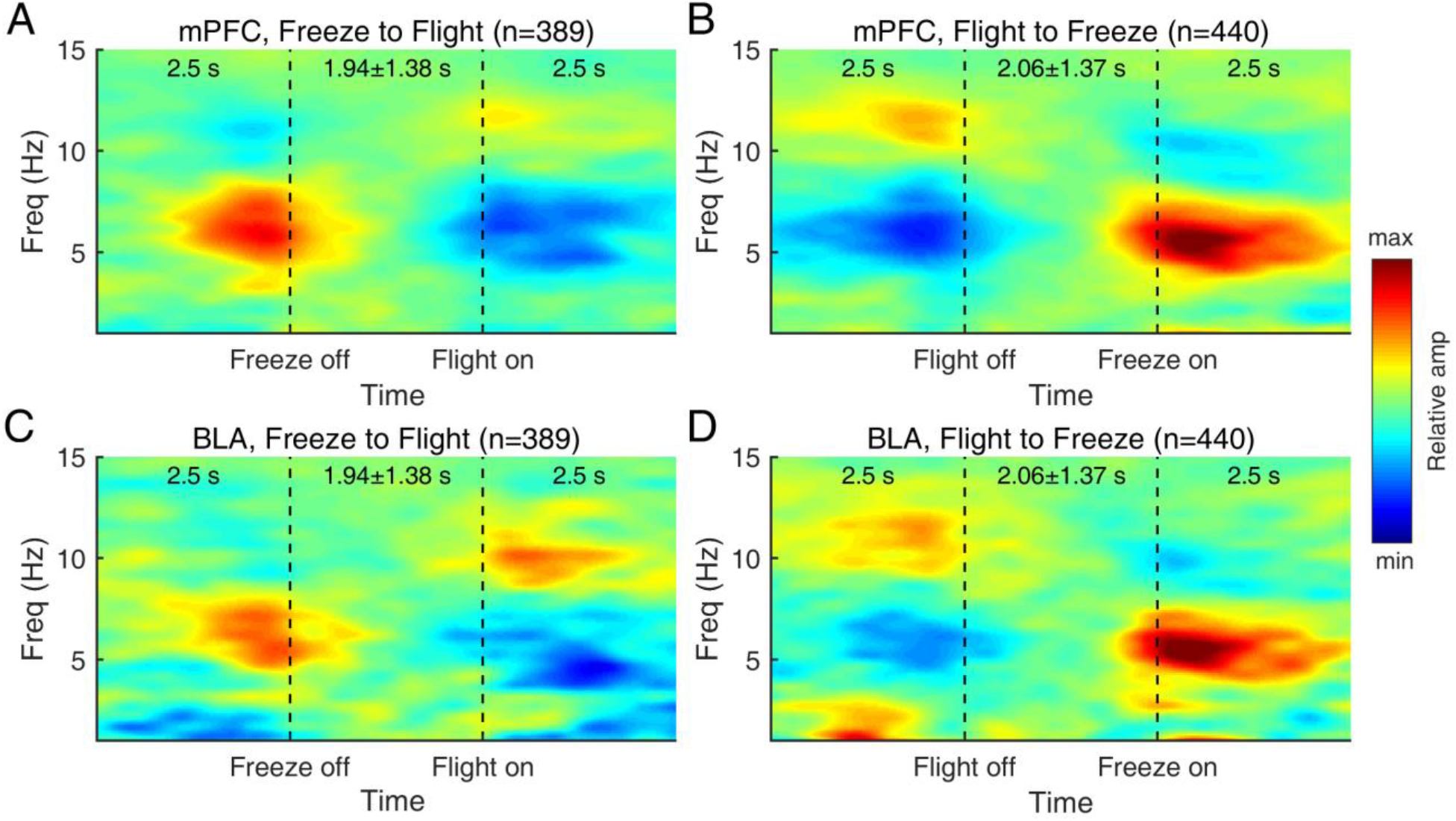
Transition between two theta rhythmic states during sequential behavioral states of freezing and flight, Related to Figure 4. (A) The animals tended to avoid the threat zone more in the final session (mean duration = 1.89 s) than in the initial session (mean duration = 17.24 s), during a time after the removal of the predator robot in Stage 4, showing their preference for the safe zone. (B) Likewise, the decrease in reaction time to the gate-opening event in Stage 3 was statistically significant (*p* = .039), showing their acquisition of task rules as well as the preference for the safe zone.

**Figure S7.**
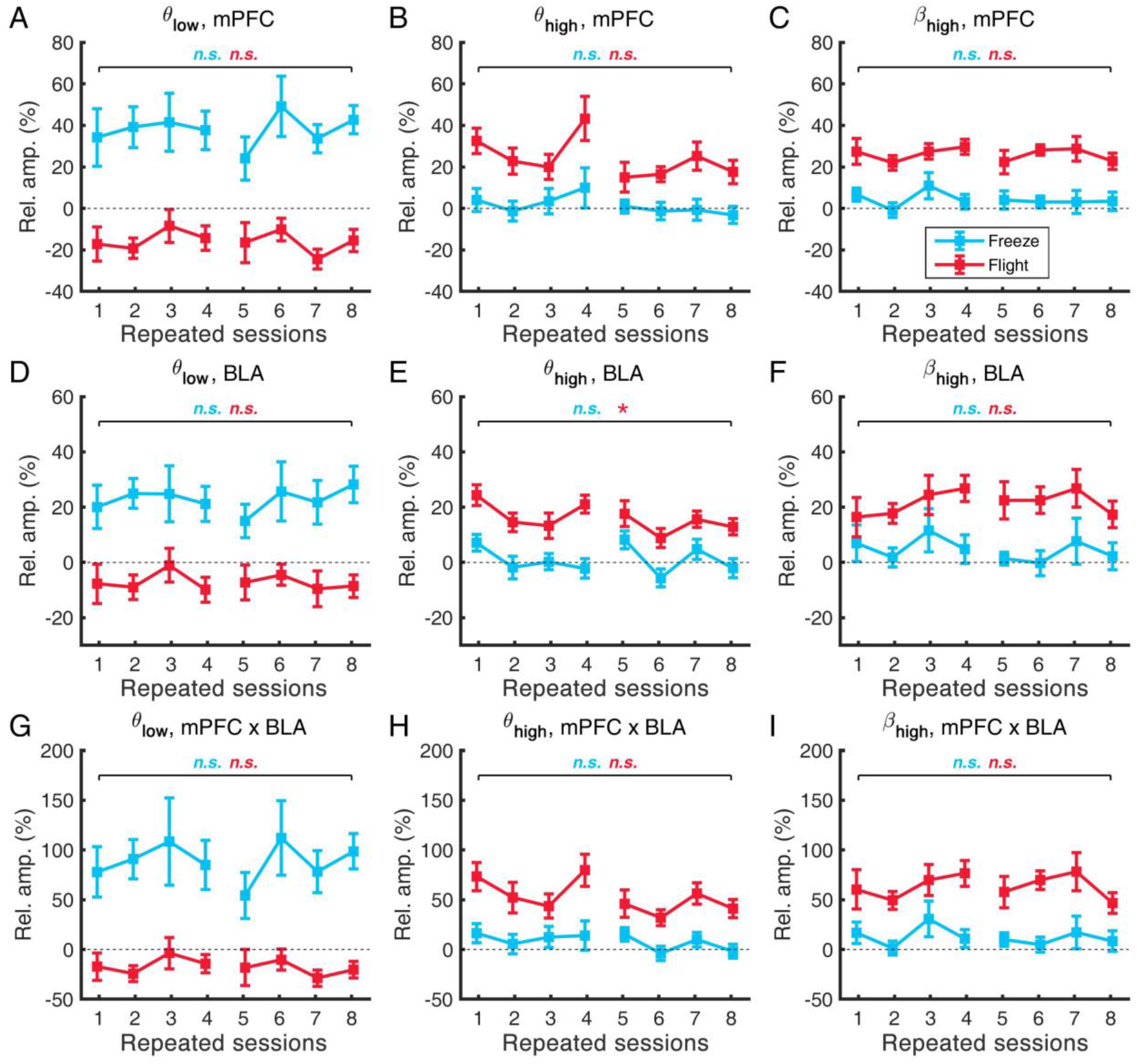
Changes across sessions in oscillatory amplitudes during two types of defensive behaviors, Related to Figure 6. (A-C). Oscillatory amplitudes in the mPFC during freezing (blue) and flight (red) states in the *θ*_low_, *θ*_high_, and *β*_high_ bands. None of the comparisons between the initial and the final sessions showed a statistically significant difference (noncorrected *p*s > .05, Wilcoxon signed rank test). (D-F). Same as (A-C), but in the BLA. Only the comparison with θ_hi_ during *flight* showed a significant difference. (G-I). Same as (A-C) but in the cross-spectra of the mPFC-BLA network. None of the pairs showed a statistically significant difference. The 1st to 4th sessions were tested on one day, and the 5th to 8th sessions were tested on the next day. * *p* < .05, ** *p* < .01. Wilcoxon signed rank test for medians of sessions 1 and 8 (α = .05, two-sided).

**Figure S8.**
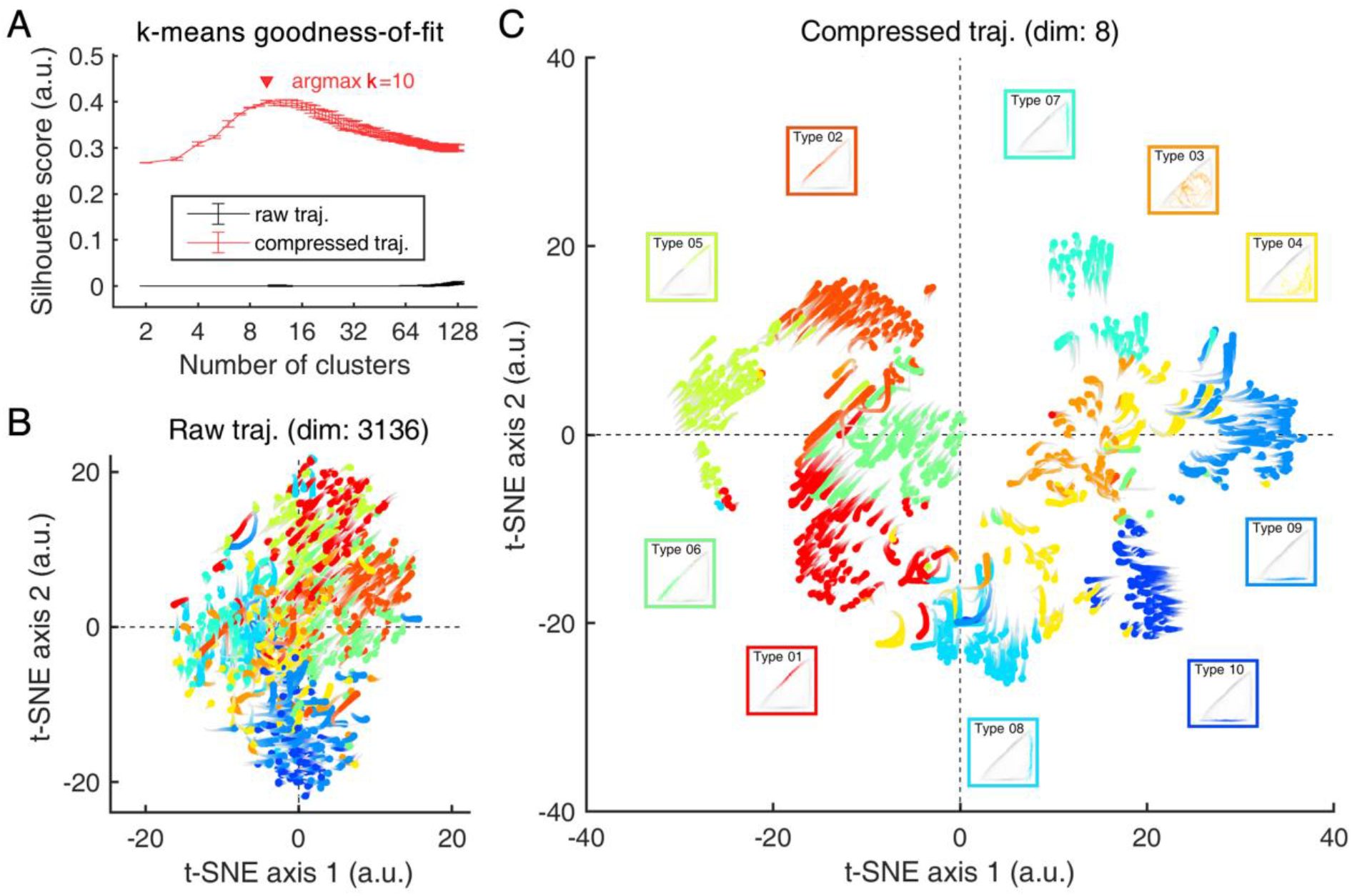
Unsupervised clustering of flight trajectories, Related to Figure 6. (A) Goodness-of-fit measured by the silhouette score to determine the proper number of clusters in k-means clustering. Error bars indicate 1 standard deviation of the mean. (B) Two-dimensional manifold embedding of raw trajectory data (dim: 56*56, binary image) performed by t-distributed stochastic neighbor embedding (t-SNE), showing poor performance of clustering due to the large dimensionality of the input. Original trajectories in the 60*60 (cm)-size arena were color-coded and rescaled in 5*5 (a.u.) for manifold visualization. (C) Same as (B) but for compressed trajectory data in the middle layer of the convolutional autoencoder. Inset images represent the grand-averaged (gray) and cluster-averaged (color-coded) trajectories.

**Movie M1.**
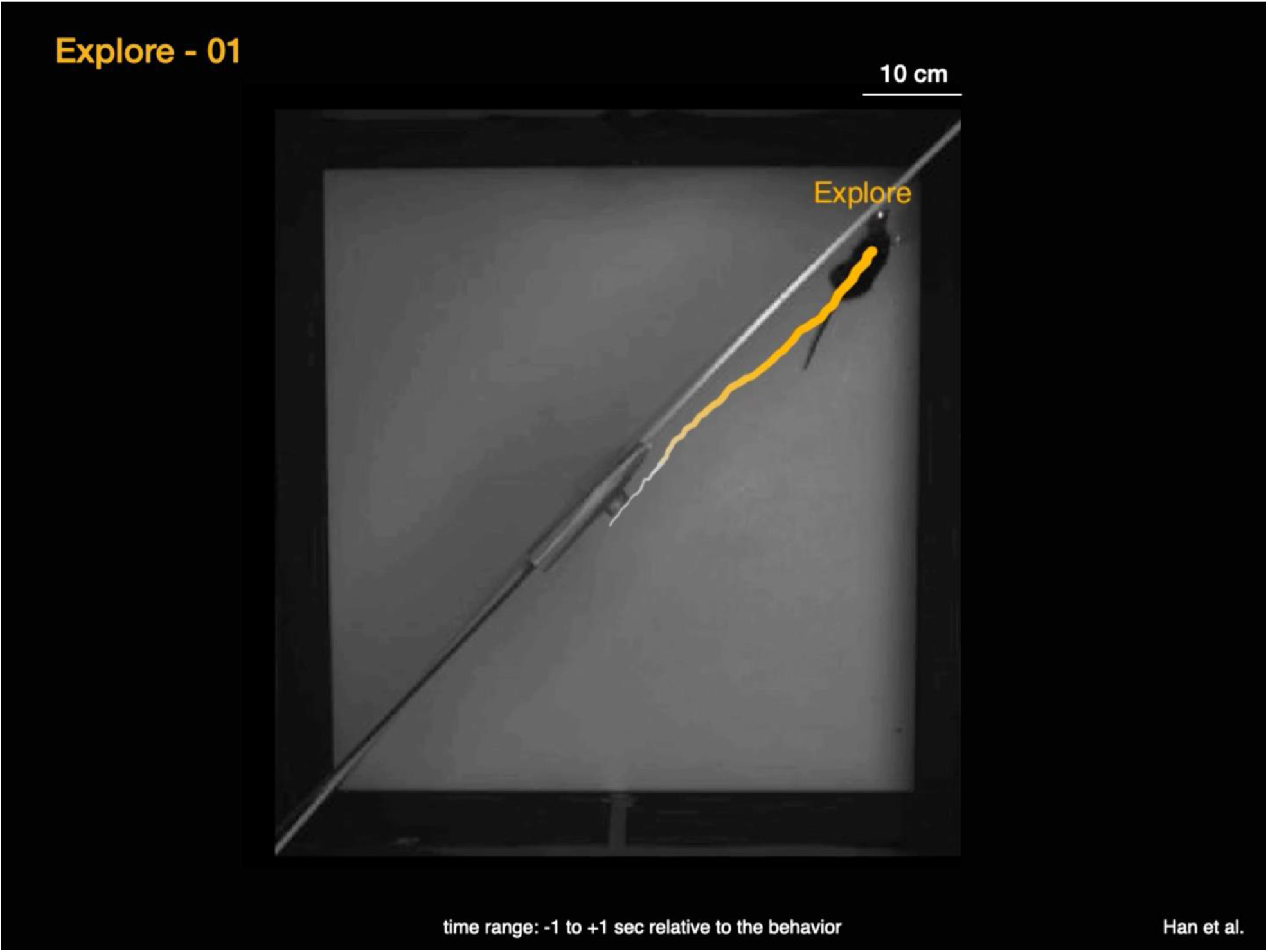
Example clips for behavioral state labeling, Related to Figure 1. Behavioral states of flight, freezing, exploring, and grooming were labeled by semi-automatic video processing. Ten (flight, freezing) or five (exploring, grooming) example bouts are shown, with the online label on the position of mouse body centroid. Thirty frames (1 s) before and after the state onset and offset, respectively, were included to demonstrate dynamic changes of behavior in our experimental setting.

**Movie M2.**
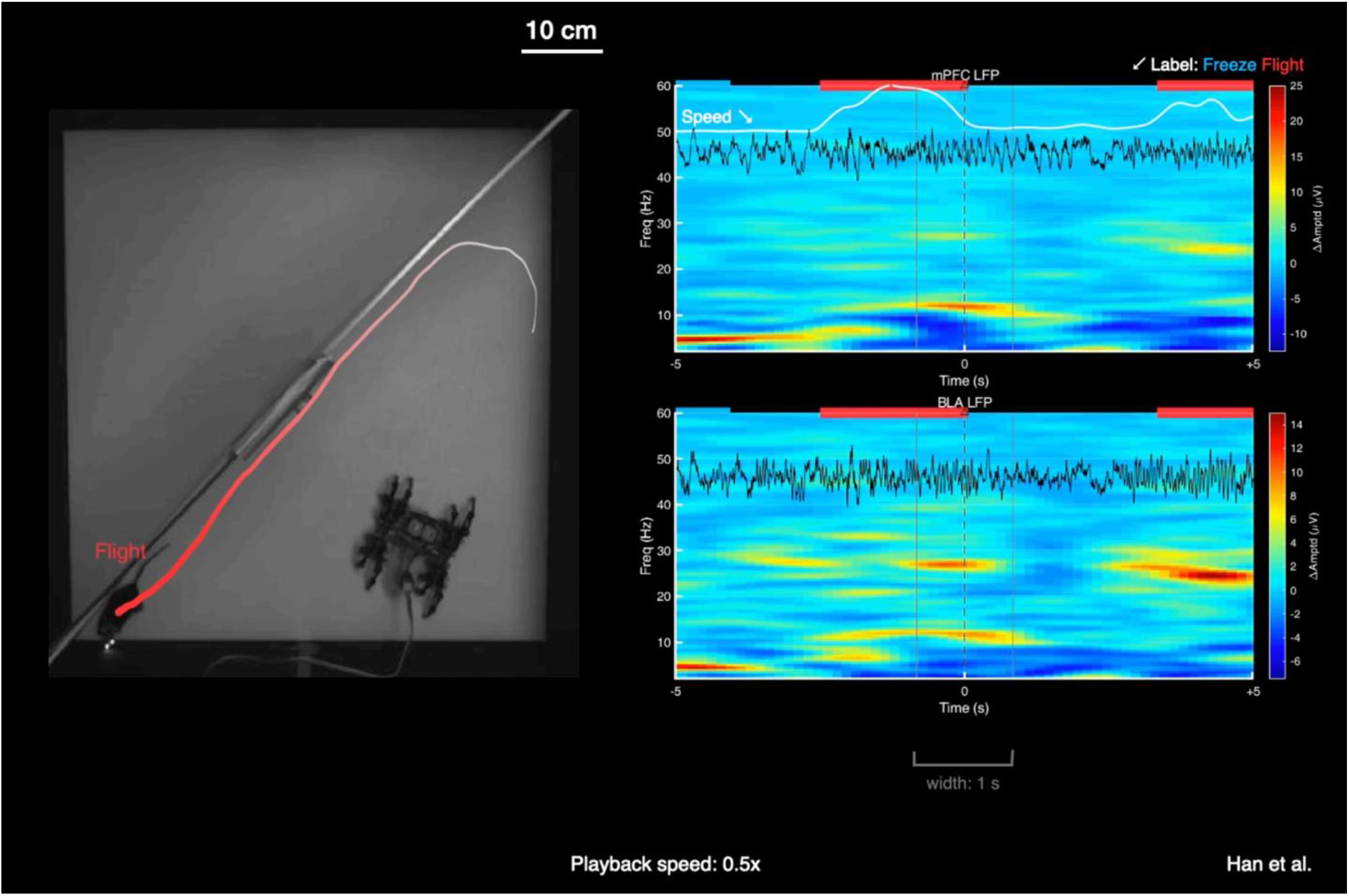
Example clip for *flight* bouts with and without mPFC-BLA beta activities, Related to Figure 6. Strong and transient beta activities in mPFC-BLA were found during flights, especially when the trajectory of flight is related to the gate of safe zone. Video playback speed is set to 0.5x.

